# *C9orf72* expansion creates the unstable folate-sensitive fragile site FRA9A

**DOI:** 10.1101/2024.10.26.620312

**Authors:** Mila Mirceta, Monika H.M. Schmidt, Natalie Shum, Tanya K. Prasolava, Bryanna Meikle, Stella Lanni, Mohiuddin Mohiuddin, Paul M. Mckeever, Ming Zhang, Minggao Liang, Ilse van der Werf, Stefaan Scheers, Patrick A. Dion, Peixiang Wang, Michael D. Wilson, Theresa Abell, Elliot A. Philips, Łukasz J. Sznajder, Maurice S. Swanson, Mustafa Mehkary, Mahreen Khan, Katsuyuki Yokoi, Christine Jung, Pieter J. de Jong, Catherine H. Freudenreich, Philip McGoldrick, Ryan K.C. Yuen, Agessandro Abrahão, Julia Keith, Lorne Zinman, Janice Robertson, Ekaterina Rogaeva, Guy A. Rouleau, R. Frank Kooy, Christopher E. Pearson

## Abstract

The hyper-unstable Chr9p21 locus, harbouring the interferon gene cluster, oncogenes and *C9orf72,* is linked to multiple diseases. *C9orf72* (GGGGCC)n expansions (*C9orf72*Exp) are associated with incompletely penetrant amyotrophic lateral sclerosis, frontotemporal dementia and autoimmune disorders. *C9orf72*Exp patients display hyperactive cGAS-STING-linked interferon immune and DNA damage responses, but the source of immuno-stimulatory or damaged DNA is unknown. Here, we show *C9orf72*Exp in pre-symptomatic and ALS-FTD patient cells and brains cause the folate- sensitive chromosomal fragile site, FRA9A. FRA9A centers on >33kb of *C9orf72* as highly-compacted chromatin embedded in an 8.2Mb fragility zone spanning 9p21, encompassing 46 genes, making FRA9A one of the largest fragile sites. *C9orf72*Exp cells show chromosomal instability, heightened global- and Chr9p-enriched sister-chromatid exchanges, truncated-Chr9s, acentric-Chr9s and Chr9-containing micronuclei, providing endogenous sources of damaged and immunostimulatory DNA. Cells from one *C9orf72*Exp patient contained highly-rearranged FRA9A-expressing Chr9 with Chr9-wide dysregulated gene expression. Somatic *C9orf72*Exp repeat instability and chromosomal fragility are sensitive to folate-deficiency. Age-dependent repeat instability, chromosomal fragility, and chromosomal instability can be transferred to CNS and peripheral tissues of transgenic *C9orf72*Exp mice, implicating *C9orf72*Exp as the source. Our results highlight unappreciated effects of *C9orf72* expansions that trigger vitamin-sensitive chromosome fragility, adding structural variations to the disease-enriched 9p21 locus, and likely elsewhere.

**GRAPHICAL ABSTRACT:** 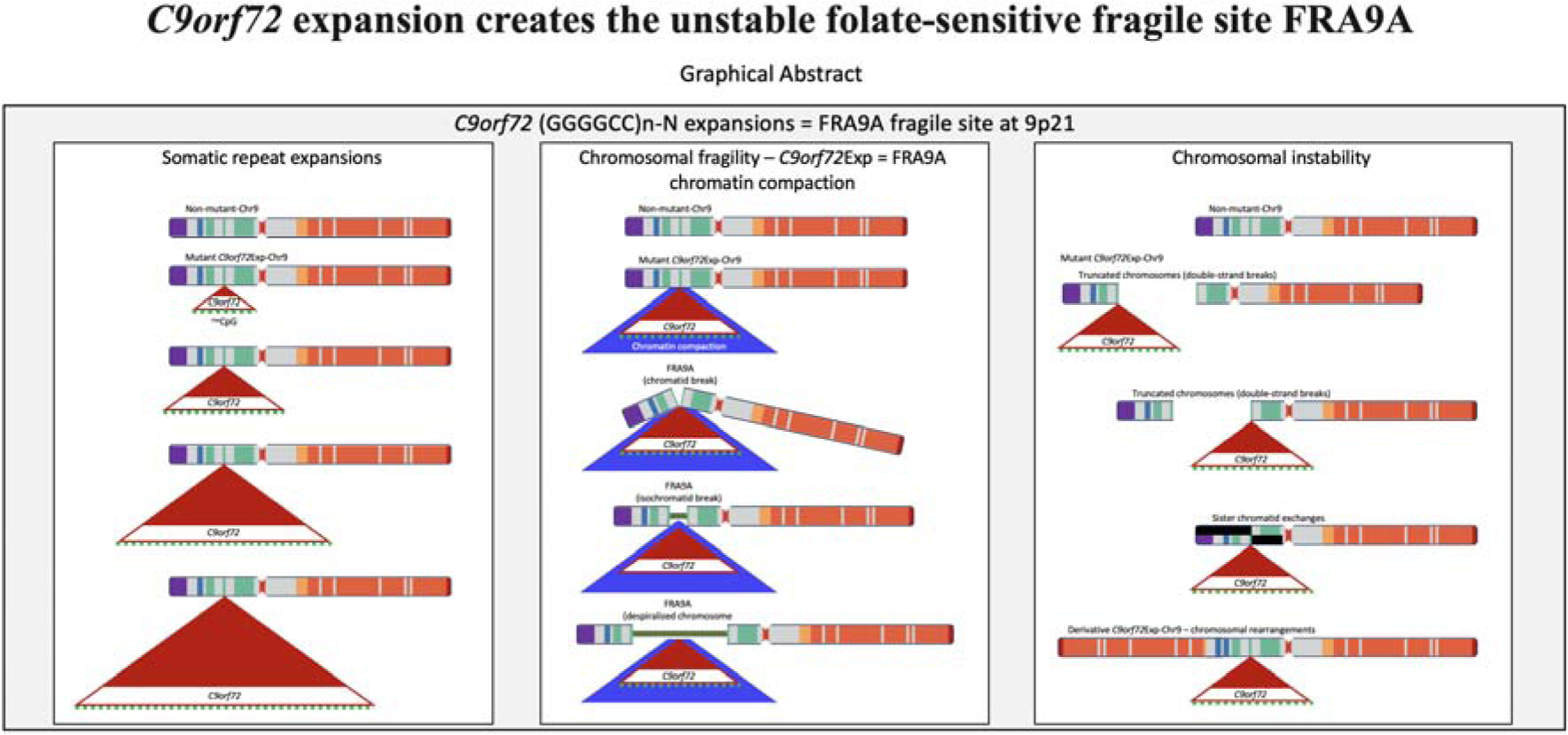

## INTRODUCTION

Expansions of 30-7950 GGGGCC repeats in the *C9orf72* gene, which resides in 9p21 (*C9orf72*Exp), are the most common cause of ALS, FTD, and numerous seemingly unrelated diseases including inflammation dysregulation, autoimmune diseases, over-expression of type I *IFN* genes (1–6), and melanoma (7). Most *C9orf72*Exp carriers are asymptomatic at 40- years of age, some will develop amyotrophic lateral sclerosis (ALS), frontotemporal dementia (FTD), another co-occurring disease, or a mixture of these, while many can be disease-free into their 80s-90s (8,9). This incomplete penetrance and clinical diversity suggests the involvement of genetic and environmental modifiers and/or cumulative events.

Many studies on *C9orf72*Exp disease have focused upon the expanded repeat RNA and repeat-associated non-AUG RAN-peptides, where many model systems are devoid of the endogenous chromosomal expanded repeat. Repeat expansions downregulate *C9orf72*Exp expression leading to reduced C9ORF72 protein levels (10, 11). *St*imulator of *In*terferon *G*enes (STING), which with cGAS activates the innate immune response following sensing of cytosolic double-stranded DNA that could arise from viruses, damaged mitochondria, or damaged chromosomes leaked from micronuclei. In *C9orf72*Exp individuals, decreased C9ORF72 leads to STING-hyperactivated *IFN* expression and inflammation in vulnerable neurons (12), but the source of the immunostimulatory DNA is unknown (4, 13, 14). Accumulating evidence supports a role of an activated DNA damage response (DDR) in *C9orf72*Exp disease (15–25), yet the source of the damaged DNA is unknown. Massive post- natal inter- and intra-tissue repeat length instability is evident in post-mortem tissues of *C9orf72*Exp individuals with the largest somatic expansions in the brain (26–30). Surprisingly, there have been no prior cytogenetic or chromosomal instability studies of *C9orf72*Exp. The recent discontinued clinical programs of antisense oligonucleotides targeting the sense-strand *C9orf72* mRNA, which successfully reduced the RAN-peptides, but resulted in no clinical benefit or trended toward greater clinical decline for the cohort, highlight our poor understanding of *C9orf72*Exp (31–36). A recent meeting focused on the post-trials path forward for *C9orf72*Exp highlighted key areas needing attention: *“Critical information in our understanding of the biology around the C9orf72 expansion is still lacking, including what molecular and cellular changes occur at presymptomatic stages…studies of genetics and lifestyles might be useful”* (31). Unknown biology of the *C9orf72*Exp has been highlighted as a focus arena (31–36). Factors that may determine disease penetrance, clinical variation, disease onset, progression and severity, are indispensable for family life decisions as well as clinical trial design (6, 9, 31, 37–41).

Genomic context is important, as *C9orf72*Exp occurs on a risk haplotype, which could influence repeat length instability, leading to reduced transcription and its retention of the intron containing the expanded GGGGCC repeat (42–45) (Fig. 1). Many of the co-occurring *C9orf72*Exp symptoms can arise independent of a *C9orf72*Exp and are associated with other 9p21 genes. For example, the type I interferon (*IFN*) cluster of 17 genes, including *IFNk* adjacent to *C9orf72*, are linked to autoimmune diseases (e.g., dysregulated inflammation, lupus, rheumatoid arthritis, diabetes, multiple sclerosis) (1, 3–5, 7). The enhancer-rich 9p21 gene desert is associated with *IFN* signaling (46, 47). *CDKN2A/B*, *EMICERI*, and *IFN*s are linked to melanoma (48), a cancer recently associated with *C9orf72*Exp (7). *LINGO2*, *TOPORS*, *APTX*, and *DNAJA1* have been associated with neurodevelopmental, neurodegenerative, and motoneuron disease (49).

**Figure 1.**
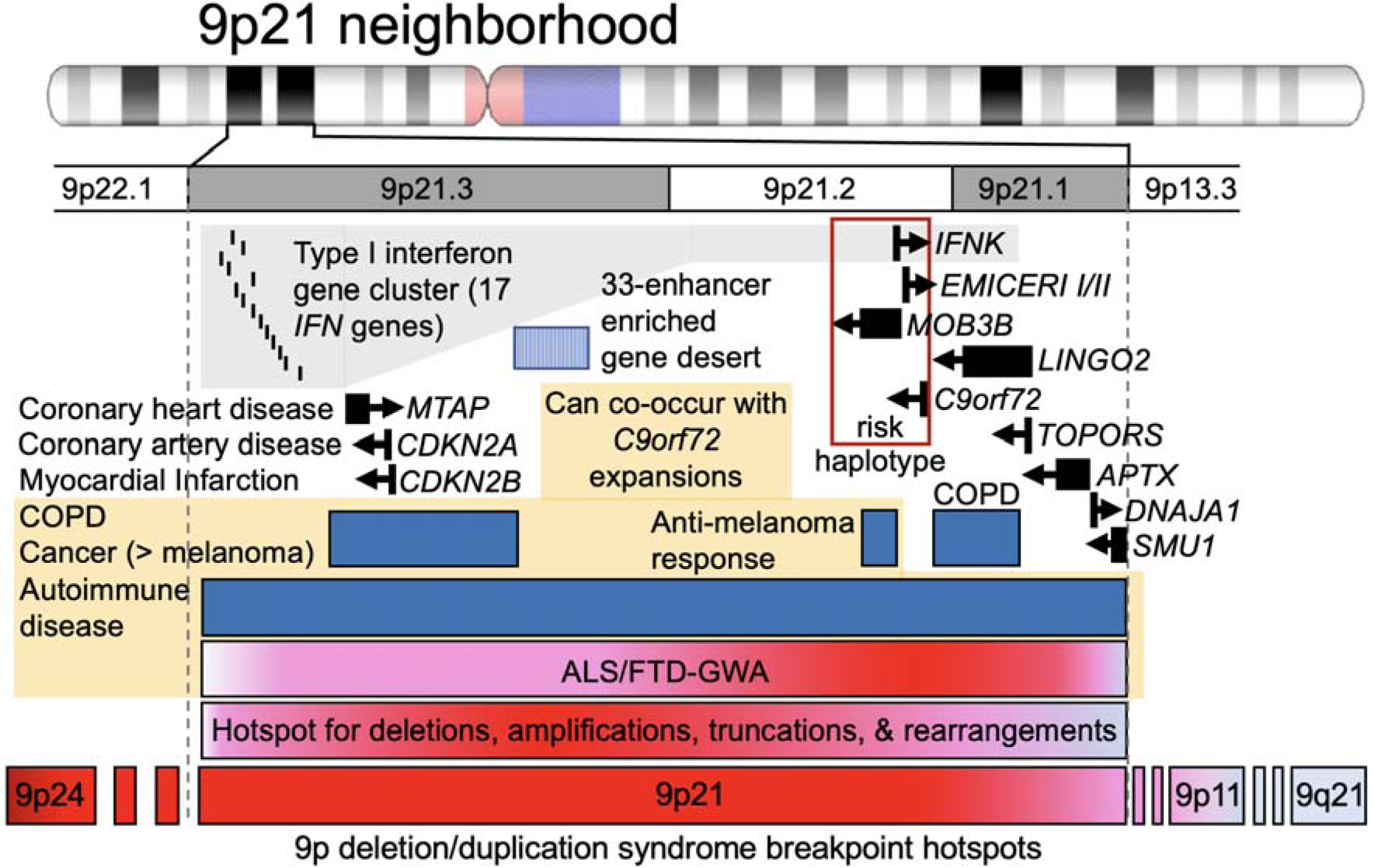
The 9p21 neighborhood, associated genes, diseases, and instability hotspots. See text.

The 9p21 locus is hyper-prone to various recurrent structural rearrangements (50–54) (Fig. 1). 9p deletion/duplication syndrome includes 9p21.2 as a breakpoint hotspot (55, 56). Instability in cancers is suspected to involve chromosomal fragile sites although the hyper-unstable 9p21 has no mapped fragile site (57). “Rare” fragile sites can be present in as few as a single individual, as initially reported for FRA9A (58), to ≤5% of individuals and are linked to partially penetrant neurological disorders, as reviewed (59). Most rare fragile sites are folate-sensitive fragile sites (FSFS), induced in the absence of folate or by the antimetabolite 5-fluorodeoxyuridine (FUdR). Ten of the ∼30 FSFS have been mapped to gene-specific expanded (CGG)n repeats, including FRAXA, FRAXE, FRAXF, FRA2A, FRA7A, FRA10A, FRA11A, FRA11B, FRA12A, and FRA16A. FSFS are prone to DNA breaks making them mutation hotspots — mutations that can alter clinical presentation, which is well-characterized for the CGG-expanded FRAXA/*FMR1*, reviewed in (59). Through >75 years of study, many isolated case reports, diverse mutation forms (varying CGG expansion sizes, contiguous gene deletions, gene duplications, intra- and inter-chromosomal recombination, point mutations), and *FMR1* mosaic epimutations have been identified as the cause of numerous seemingly unrelated diseases, many incompletely penetrant (59). Individual case studies with deep clinical, neuropathological and molecular phenotyping can be informative of disease etiology. Thus, instability of FSFS can impact disease manifestation and clinical presentation. Our objective was to better understand the biology of the endogenous expanded *C9orf72* repeat in the context of 9p21 in presymptomatic and disease states. We assessed whether the expanded *C9orf72* repeat, GGGGC***CGG***GGC***CGG***GGC***CGG***GGCC…, which contains the CGG-motif common to all mapped FSFS, is a FSFS.

## RESULTS

### 9p21 shares sequence and epigenetic features with (CGG)n FSFS

The *C9orf72* locus and each of the CGG-expanded FSFS share many sequence, structural, functional, and epigenetic features (Fig. 2A-D, Fig. S1A-J). Nine of the 10 CGG-FSFS plus *C9orf72* colocalize to boundaries between topologically associating domains and were enriched with CTCF sites and CpG-islands (Fig. 2D). Thus, *C9orf72* and CGG-FSFS share with 22 other disease-associated repeat expansion loci these epigenetic features (60). Unlike other disease-associated repeats, both the *C9orf72* GGGGCC and all CGG-FSFS share G4- quadruplex forming repeats (61–63).

**Figure 2.**
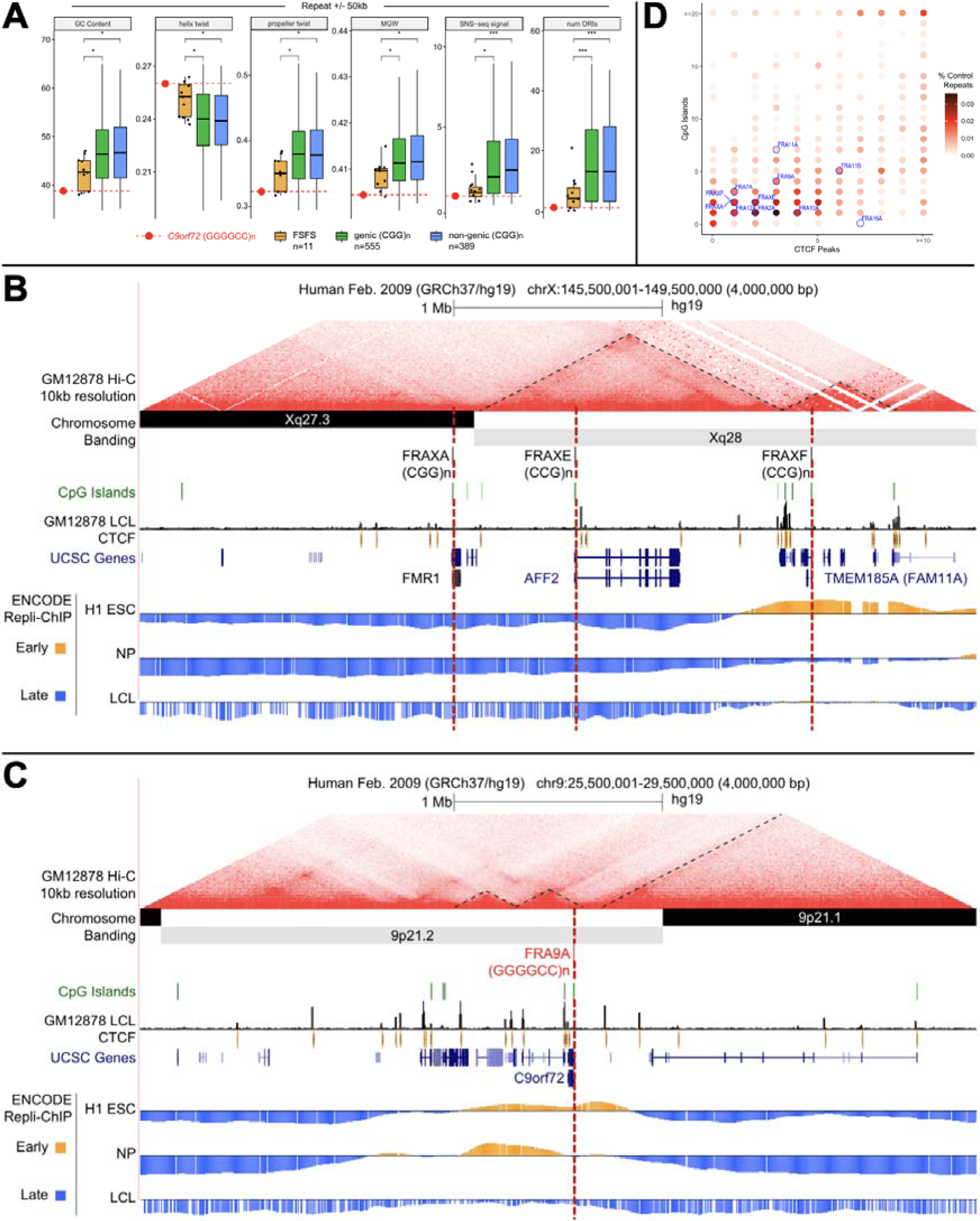
DNA sequence, structure, topological associated domain (TAD), CpG island, and Repli-Chip analysis for all 10 CGG-folate-sensitive fragile sites. (A) Boxplots comparing average %GC, helix twist, propeller twist, major groove width (MGW), SNS-seq (short nascent strand sequencing) signal and number of origins of replication initiation (ORIs) at CGG/CCG-mapped FSFS and *C9orf72* versus 944 control genome-wide CGG/GGC repeats, calculated for windows +/- 50kb and +/- 500kb from the repeat center. Control repeat regions were further separated as genic or non-genic based on overlap with an annotated transcript. Median repeat length for FSFS is 8.7; all FSFS have GC content of 100% (AT=0 except FRAXF, GC=0.826). Repeat length req of >4.5 for 944 CGG/CCG control regions yieded a median length of 8.7, same as mapped FSFS. Asterix denotes significance based on a two-sided Wilcoxon test (* p<0.05; ** p<0.01). Each individual FSFS is represented as a black dot. The *C9orf72* (GGGGCC)n is represented by a red dot. (B & C) Topological domain, DNA replication timing tracks, CpG islands, and CTCF sites for the contiguous FRAXA, FRAXE, and FRAXF compared to same for *C9orf72*. Shown are the contour density plots depicting the number of CTCF sites and CpG-islands in 100 kb windows centered on the repeat, representing boundaries with normal-length, matched repeats. Repli-seq data were obtained in bigwig format (GSE61972). H1 ESC = H1 Embryonic Stem Cells; accession:ENCFF000KUF; NP = Neuronal progenitor cells (BG01 Fibroblast-derived); accession:ENCFF907YXU; LCL = GM06990 lymphoblastoid cells; accession:ENCFF000KUA. Early (positive) and late (negative) replication was determined by subtracting the genome-wide median score from all values. To conserve space, only the FSFS gene is indicated, while other UCSC genes are condensed into one line. (D) Summary of enrichment of each fragile site (indicated in blue fonts) for CpG islands and CTCF sites. Points are colored according to density. For analyses of FRA2A, FRA7A, FRA10A, FRA11A, FRA11B, FRA12A, and FRA16A see Fig. S1.

### Variations in *C9orf72* repeat expansion size, CpG-methylation, and disease state permit assessment of contribution to fragility

To study the connection between *C9orf72*Exp and fragility, we characterized lymphoblastoid cells from *C9orf72*Exp families, where some carriers were presymptomatic and some presented with ALS or ALS/FTD (Fig. 3A). Sizing *C9orf72* expansions by Southern blots shows repeat length heterogeneity within and between individuals, with expansions ranging from ∼300 to ∼4400 repeats (Fig. 3B, Table S1). All individuals carried (GGGGCC)2 on the non-expanded allele.

**Figure 3.**
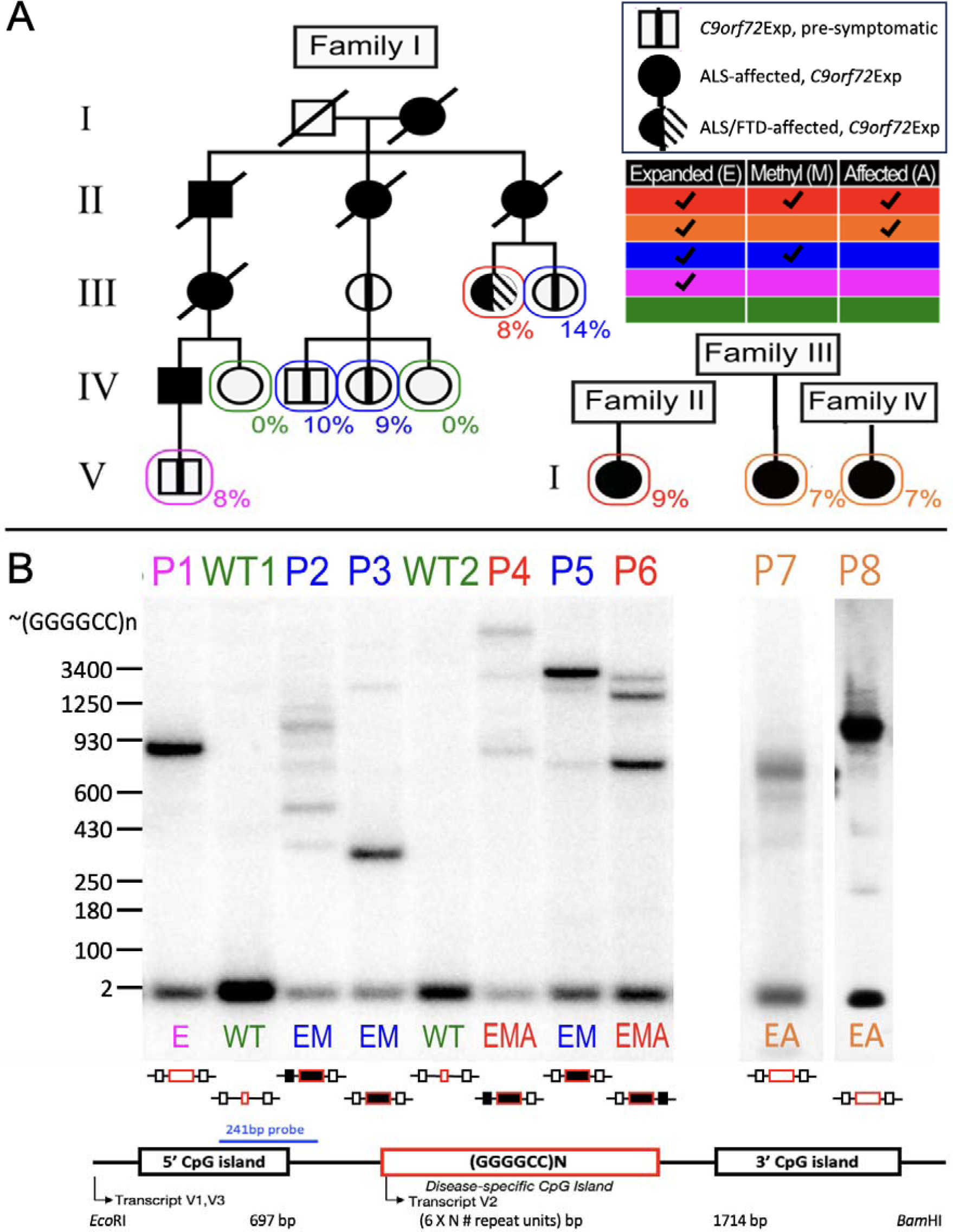
*C9orf72*Exp cells, repeat lengths, CpG-methylation, and disease state. (A) ALS/FTD families with Southern blot vertically aligned for each individual. Females are represented by circles, males by squares, disease state is as per the insert legend. Colors of text and boxes around symbols indicate the expansion, methylation, and affected status for each sample. Percentage of cells showing FRA9A expression are indicated for each individual in the pedigree. “P#” refers to patient number, “E”, “M”, and “A” refer to *E*xpanded, *M*ethylated, and *A*ffected, is summarized at right and this coding system is used throughout the text. Southern blot of *C9orf72* (GGGGCC)n repeat expansion (Methods) used *Eco*RI/*Bam*HI to release the repeat-containing fragment leaving 697 and 1714 bp upstream and downstream of the (GGGGCC)n/N repeat. (B) Schematic of Southern blot probe location, CpG-methylation status at the GGGGCC repeat and CpG-islands (filled boxes denote methylation). For raw CpG methylation data see Figure S2.

Most of our cell lines were CpG methylated in the expanded repeat, with varying levels in flanking CpG-islands, and no methylation in the non-expanded allele (Fig. 3B, Fig. S2), consistent with previous findings of aberrant methylation and levels of methylation mosaicism in *C9orf72*Exp carriers (64, 65). We found an absence of methylation, likely arising from methylation mosaicism, in cells of a presymptomatic *C9orf72*Exp (P1) and ALS-affected individuals (P7 & P8). No obvious differences were observed between cells from presymptomatic and affected *C9orf72*Exp individuals. Methylation status was stable, consistent with longitudinally collected samples from *C9orf72*Exp carriers (66). Control cells from individuals without *C9orf72*Exp had no methylation at *C9orf72*. Thus, our *C9orf72*Exp cell line collection permitted assessing fragility in the presence/absence of methylation and with/without disease of donors (Fig. 3A, see right-hand chart).

### *C9orf72* expansion is the molecular cause of a FSFS at 9p21 (FRA9A)

Fragility was assessed cytogenetically (Fig. 4A-D). FUdR-induced fragile sites were observed and localized to 9p21 by fluorescent *in situ* hybridization (FISH), in all eight *C9orf72*Exp cell lines. Fragility was only observed in *C9orf72*Exp cells, with FUdR- treatment and only on one Chr9, consistent with FRA9A (58). Fragility presented in various forms including gaps (isochromatid and chromatid), constrictions, and despiralized regions (Fig. 4A). Full metaphase spreads and zoomed-in versions, with greater resolution are available upon request. With the exception of the despiralized form, these are similar to forms of FRAXA (59). The despiralized Chr9 was similar to the despiralized heterochromatin in immunodeficiency, centromeric instability, and facial anomalies (ICF) syndrome (67). Lengthening of the despiralized FRA9A is evident when compared to the non-fragile Chr9 from the same metaphase spread (Fig. 4A, control Chr9 in lower left is from the same metaphase as the leftmost stretched FRA9A). These cytogenetic observations reveal a fragile site, FRA9A, at 9p21.

**Figure 4.**
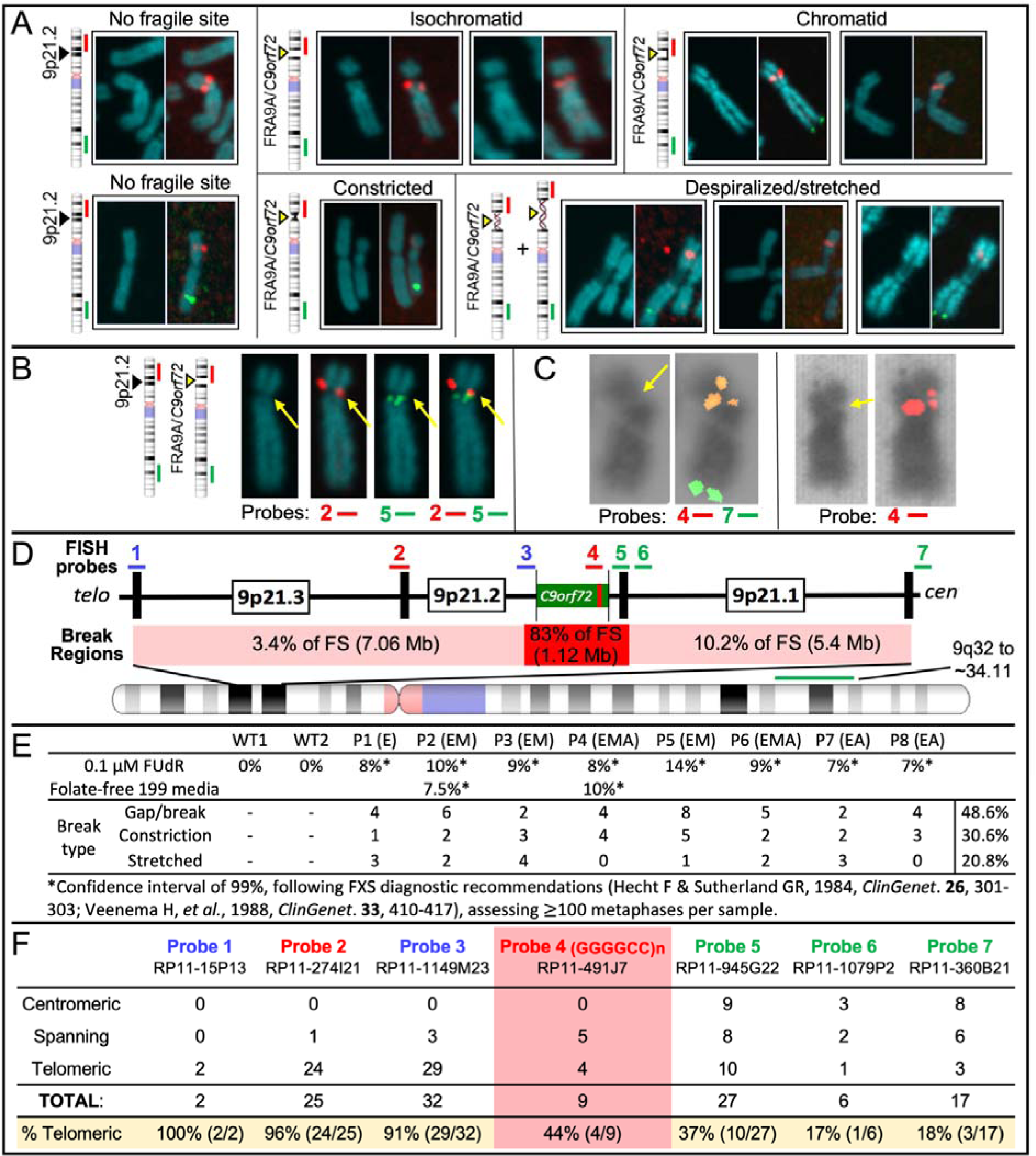
*C9orf72*Exp causes FRA9A folate-sensitive fragile site that maps across 9p21. (A) Fragile site forms (indicated), with examples of non-fragile chromosomes from the same metaphase spreads. Showing DAPI-stained and paired FISH-probed chromosomes from FUdR-treated cells. 9q-arm FISH probe (9q32∼34.11; RP11-696J10) in green. FISH probe #2 at *C9orf72* (panel D- F) in red. Magnification is the same for all chromosome images within each panel, facilitating comparison within the panel. Control Chr9 in upper left of panel A is from the same metaphase as the leftmost isochromatid Chr9. Full metaphase spreads can be provided upon request. Notably, the control “No fragile site” Chr9 in lower left of panel A is from the same metaphase as the leftmost “Despiralized/stretched” Chr9, revealing the lengthening of the stretched Chr9. (B) Representative example of the same chromosome (same magnification) with or without indicated FISH probes. (C) Two representative examples (same magnification) of isochromatids with FISH signal of *C9orf72*- containing probe 4 split over the fragile site. (D) 9p21 location of FISH probes with regional summary of fragile site location. (E) Quantification of fragility for 100 metaphases under two folate-stressed conditions. (D) Cells were described in Figure 3. (F) Fragile site breakage regions calculated from number of breaks relative to FISH signal found centromeric, telomeric, or spanning across probes used. Percentage of telomerically located FISH signal relative to the break is on the bottom row. Frequencies of probes found spanning or on each side the break were used to calculate the percent breakage within each region in panel D.

We next quantified fragility. Established FXS/FRAXA diagnostic guidelines (68, 69) dictate that fragile site frequencies of 3% or 5% within 100 metaphases yields confidence levels of 95% or 99%. Typically, FXS individuals show fragility at 3-15%, but never >50% of CGG expanded ChrXs (69, 70). We observed 9p21 fragility at 7-14% across *C9orf72*Exp cell lines (Fig. 4E). Fragility was evident in *C9orf72*Exp cells of presymptomatic and ALS/FTD-affected individuals. Chromosomal assignment and band location of fragility was confirmed by Giemsa-banding, trypsinization, and karyotyping (Fig. S3). Fragility at 9p21 was observed on only one of the Chr9 homologs in a given metaphase of any one *C9orf72*Exp cell line, consistent with each being heterozygously expanded. Fragility at 9p21 was only detected with FUdR treatment. Similar levels of fragility arose using folate-free media (M199) as for FUdR, revealing induction by different methods of folate stress (Fig. S3A). Fragility could not be induced without *C9orf72* expansions in two control cell lines (100 metaphases each). Thus, fragility depends on folate perturbation and the presence of the expanded repeat. As few as 300 repeats (P3, 9%) could express FRA9A at similar levels as 4400 repeats (P4, 8%). The localized fragility, limited to one Chr9, in *C9orf72*Exp cells is consistent with previous FS mapping to autosomes that concluded fragility arises on the expanded allele (71, 72). We validated that the *C9orf72* repeat expansion is the molecular cause of FRA9A fragility through gene-transfer experiments (below).

We further mapped fragility using seven FISH probes spanning the outer edges of 9p21, in eight *C9orf72*Exp cell lines, >100 metaphases each, with probe signals being telomeric, spanning, or centromeric to the break (Fig. 4B, 4D & 4F). Breaks were detected across 9p21.1-9p21.3 chromosomal bands, ∼8.2Mb, with ≥83% of breaks at a ∼1Mb stretch of 9p21.2 encompassing *C9orf72* (Fig. 4D). Splitting of the FISH signal to both sides of the break was evident for the GGGGCC-containing probe 4, supporting that the expanded repeat causes fragility (Fig. 4C). Probes telomeric to *C9orf72* preferentially fluoresced telomeric of the break, with the same trend for centromeric probes (Fig. 4F). The telomeric boundary could be considered at probe 2, which straddles 9p21.2-9p21.3, with fluorescence both telomeric and spanning fragility. A centromeric boundary could not be defined, as probe 7 was detected telomeric, spanning, and centromeric to breaks, suggesting fragility extended centromeric of probe 7. Additional centromeric probes, being pericentromeric, hampered break detection. Thus, within the limits of FISH resolution, FRA9A maps most intensely to *C9orf72*, but has a wide zone of breakage (summarized in Fig. 4D). FRA9A’s fragility core of enriched breakage includes *C9orf72* covering ∼500kb (plus the repeat), spanning the *C9orf72*Exp risk haplotype, where breaks can extend ∼2Mb and ∼6.2Mb telomerically and centromerically (Table 1). This makes FRA9A one of the largest fragile sites, spanning

**Table 1:** Large fragile sites with variably located fragility cores (increased breakage) and gene numbers.

| Fragile site | location | fragility size | fragility core size | core location | # genes* | TEL schematic; — = 1 Mb CEN | citation |
| --- | --- | --- | --- | --- | --- | --- | --- |
| FRA1H | 1q41-42.1 | 10 Mb | 4 Mb | asymmetric | 49 |  | 75 |
| FRA3B | 3p14.1-14.2 | 4 Mb | 500 kb | asymmetric | 24 |  | 77 |
| FRA4F | 4q22 | 7 Mb | 4 Mb | broad | 12 |  | 76 |
| FRA7B | 7p21.3-22.3 | 12.2 Mb | 5 Mb | asymmetric | 79 |  | 73 |
| FRA9E | 9q32-33 | 9.8 Mb | 5 Mb | broad | 63 |  | 74 |
| FRA9A | 9p21 | 8.2 Mb + repeats** | 500 kb + repeats** | asymmetric | 46 |  | this study |
| FRA11G | 11q23.3 | 4.5 Mb | 2.32, 1.37 Mb | broad | 38 |  | 78 |
| FRAXA | Xq27.3 | 20-600 kb + repeats** | 20 kb + repeats** | centric | 10 |  | 79, 80 |
| FRAXF | Xq27-q28 | 1-2.5 Mb + repeats** | 1 Mb + repeats** | asymmetric | 110 |  | 81, 82 |
| *The number of genes in the fragile region was determined for protein-coding genes as of 01-2024 as per FISH probes used.<br>**Schematic of FRA9A, FRAXA, and FRAXF is shown with (GGGGCC)4000, (CGG)900, and (CCG)600 repeats, respectively (red bars). |  |  |  |  |  |  | fragile<br>fragility core<br>dispersed fragility<br>repeat |

∼8.2Mb over 9p21.1-21.3, encompassing 46 genes (Table 1), similar in size to FRA1H, FRA3B, FRA4F, FRA7B, FRA9E, FRA11G, and FRAXF, spanning up to 12.2Mb over multiple chromosome bands, encompassing 10-110 genes (Table 1) (73–82). Moreover, CpG-methylation of *C9orf72*Exp was not required, as fragility was observed in cells +/- methylation (Fig. 4E). Thus, neither symptomatic state of blood donors nor CpG-methylation were required for FRA9A expression. That we detected FRA9A fragility at 9p21 in *C9orf72*Exp, lymphoblastoid cells, confirms previous observations that FRAXA expression was similarly detected lymphoblastoid cells as in freshly-collected peripheral blood lymphocyte cultures (83). Thus, FRA9A and FRAXA are distinct from the fragility at 11q23, which depends upon the EBV encoded protein EBNA1 (84–87).

### *C9orf72*Exp allele assumes a compact chromatin conformation, which is enhanced by expansion size and CpG-methylation

Since constricted fragile sites on metaphase chromosomes may be the result of localized unusual chromatin condensation, we next determined whether the mutant *C9orf72*Exp locus assumes an unusual chromatinized state relative to the non-mutant allele using micrococcal nuclease (MNase) accessibility, which is widely used as an indication of regional chromatin compaction. We were able to electrophoretically resolve MNase accessibility on the mutant from the non-mutant alleles by Southern blotting. MNase can directly reveal both inaccessible and accessible regions for each allele, while sequencing-based methods of chromatin accessibility, like ATAC-seq, must impute inaccessibility and cannot resolve alleles. Permeabilized cells were exposed to MNase, which preferentially cuts DNA between nucleosomes; DNA was de-proteinized, isolated, restriction digested releasing the repeat-containing fragment, electrophoretically resolved, and detected by Southern blotting (Fig. 5A-B, blue probe). Over-digestion by MNase to mono-, di-, and tri-nucleosome sized DNA fragments was evident by ethidium staining (Fig. 5C, left panel). A striking resistance to MNase accessibility is seen exclusively for the *C9orf72*Exp allele, contrasting the non-expanded allele, which serves as an internal equimolar control, completely degraded at low MNase concentrations (Fig. 5C, compare lanes 4 & 5). The expanded allele resisted digestion at much higher MNase concentrations (Fig. 5, compare lanes 5-10). MNase resistance extended beyond the expansion into the flanking regions of the mutant allele. Increasing concentrations of MNase (>50 Units) led to partial and progressive degradation of the expanded allele to a distinct shorter size, losing ∼1 kb of flanks from the full-length *Eco*RI- *Bam*HI restriction fragment, which subsequently digested the probed region (Fig. 5C, schematic). Conversely, on the non-mutant allele, the same flanking regions and non-expanded repeat were completely digested at low MNase concentrations, along with the rest of the genome (<10 Units, Fig. 5C, lanes 5-10). The poor digestion of the *C9orf72*Exp mutant allele relative to the internal sequence control of the equimolar non-mutant allele, strongly suggests the formation of an unusual chromatinized state of the mutant *C9orf72* allele. Control experiments confirmed that the inaccessibility of MNase to the *C9orf72*Exp allele was due to its chromatinized state, and not due to an inherent inability of MNase to cleave the expanded non-chromatinized (GGGGCC)n sequence (Fig. S4A & S4B, Methods). Six cell lines showed MNase resistance of the *C9orf72*Exp allele, regardless of presymptomatic or symptomatic state of the blood donors (Fig. 5C-E).

**Figure 5.**
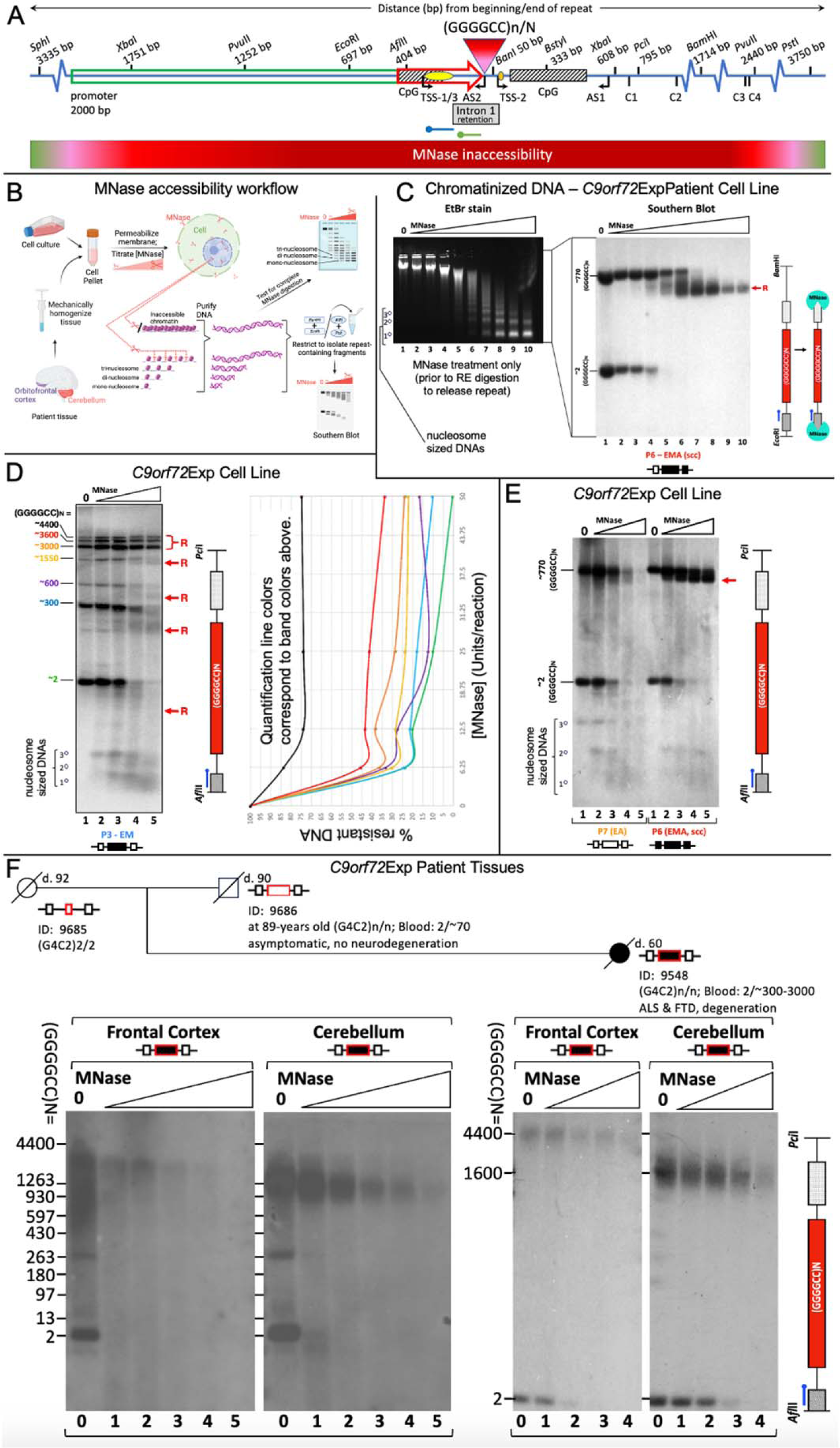
C***9***orf72Exp **locus is MNase-inaccessible.** Schematic of the region of unusual chromatin organized as determined by MNase accessibility analysis in *C9orf72*-ALS patient cells and brain regions. *C9orf72* (GGGGCC)n-N repeat, CpG-islands (hatched boxes), long-2kb and intermediate 435bp promoters (green and red arrows, respectively), exons 1a (yellow dot) and 1b (orange dot), sense and antisense transcription start sites, restriction sites with distances from the beginning and end of the (GGGGCC)N tract, and probes for upstream (green) and downstream (blue) mapping of MNase accessibility. Cryptic splice site transcripts C1-C4 derived from the *C9orf72*Exp allele are indicated with their distance from the end of the repeat tract (42, 45). Workflow for MNase assay. (C) MNase treatment of chromatinized DNA from a cell line (P6) with a large methylated *C9orf72*Exp that had undergone single-cell cloning (scc) to isolate a single expanded allele, ensuring a single expansion size and facilitating clarity. Ethidium bromide (EtBr) staining of the MNase-treated DNA reveals over-digestion of the DNA to mono-, di-, tri-nucleosomal, sized fragments. The same DNA was digested with *Eco*RI/*Bam*HI, to release the repeat fragment having 697 bp upstream and 1714 bp downstream of the repeat, then assessed by Southern blotting using the blue probe. Resistance of only the expanded allele occurs, even at high concentration of MNase (200U). Red “R” indicates the region with reduced MNase accessibility that is about to lose the probed region indicated in the schematic at right. (D) Sample P3 with a mixture of heterogeneous repeat allele lengths shows that with increasing repeat size, MNase accessibility is decreased, as quantified by densitometric analysis for each allele size (colours coordinate between graph and Southern blot). In panels D-F the repeat was released with *Afl*II/*Pci*I, leaving 404 bp upstream and 689 bp downstream allowing increased resolution. NB, van Blitterswijk uses *Xba*I/*Xba*I to release the repeat, leaving 1751 bp upstream and 608 bp downstream of the repeat, the greater amount of flanking sequence reduces electrophoretic resolution. (E) Methylation is associated with increased MNase inaccessibility as assessed in cell lines (P6 (ssc) and P7) containing similar sized repeats but differing methylation status. The non-expanded allele is digested at the same rate in both lines. (F) MNase treatment of post-mortem (orbito)frontal cortex and cerebellum from an asymptomatic 90- year-old father and his ALS-FTD-affected daughter, previously deep-phenotyped for lifestyle choices, clinical, neuropathological, and molecular biomarkers (29, 88) (summarized in Table S2). MNase concentrations are doubled between each lane (2.5U in lane 2 up to 40U). Expanded allele in both tissues is MNase resistant relative to the equimolar non-expanded allele, which serves as an internal control. See Figure S4 for MNase controls and mapping experiments.

Mapping the boundaries of MNase-resistance using multi-probe and multi-restriction digests revealed the extent and distribution of MNase inaccessibility over the *C9orf72*Exp allele. Maximal MNase resistance spanned a total of >33kb for an allele with 4400 repeats (P4), with MNase inaccessibility extending ∼3300bp upstream and ∼3700bp downstream of the repeat (summarized in Fig. 5A to scale, Fig. S4C, see Methods). Mapping in six *C9orf72*Exp cell lines showed little variation of the absolute boundaries of MNase inaccessibility with expansion size or presence of disease. Together, these data confirm that the *C9orf72*Exp allele assumes a large region of compact chromatin. Maximal MNase inaccessibility covers the upstream CpG-island, the *C9orf72* promoter, transcription start sites (sense and anti-sense), the expanded repeat, cryptic splice sites C1-4, the downstream CpG-island, beyond exon 2, and falls within FRA9A’s fragility core.

Repeat length and CpG-methylation enhanced MNase-resistance. Analysis of a cell line that harbored a mosaic heterogenous mixture of expansion sizes ranging between ∼300–4400 repeat units demonstrated that repeat expansion size directly correlates with MNase resistance (Fig. 5D, P4). Longer expansions were more resistant to MNase than shorter expansions, which were more resistant than the non-expanded alleles (Fig. 5D, lanes 3 & 4, quantified in Fig. 5D). CpG-methylation exacerbated inaccessibility. Two cell lines with similar expansion sizes (∼770 repeats) with or without aberrant methylation at the mutant *C9orf72* (P6 [isolated single cell clone] and P7, respectively) showed that methylation further enhanced MNase resistance of the expanded allele (Fig. 5E, S4A-iii). Larger expansions and methylation enhance chromatin compaction of the *C9orf72* locus.

We assessed chromatin compaction of post-mortem brains of two *C9orf72*Exp individuals, previously deep-phenotyped for lifestyle choices, clinical, neuropathological, molecular and genetic biomarkers (29, 88). One was a 90-year old male asymptomatic for ALS or FTD, with a distinct ∼70 repeats in blood, no signs of neurodegeneration, overexpression of *C9orf72*/C9ORF72, and devoid of TDP-43 neuropathology (summarized in Table S2). The other was his daughter with 300-3000 repeats in blood who had ALS at age 57 and died at 59 and autopsy showing neurodegeneration and TDP-43 inclusions (29) (detailed in Table S2). The asymptomatic father had a striking heterogenous mosaic range of 13-3500 and 13-1500 repeats in the frontal cortex and cerebellum, respectively, with CpG-methylation likely triggered by large (>70 repeats) expansions (Fig. 5F, lanes 0). In contrast, the affected daughter showed very large distinct expansions of mostly ∼4400 and ∼1600 CpG-methylated repeats in the frontal cortex and cerebellum, respectively, where the intensity of the expansions matched that of the non-expanded allele (Fig. 5F, lanes 0). Chromatin was MNase inaccessible on only the expanded *C9orf72* alleles in both brain regions of both individuals, while the non-expanded (GGGGCC)2 allele was rapidly digested by MNase (Fig. 5F). MNase inaccessibility for both individuals appeared more severe in the cerebellum relative to the frontal cortex, which contrasts the expectation that the longer expansions in the frontal cortex would make it more MNase resistant, but may reflect the longer expansions and/or the increased levels of deacetylated and trimethylated histone modifications in the frontal cortex over the cerebellum, relative to the acetylated histones on the non-expanded allele of *C9orf72*Exp carriers (89). The shorter expansions in the father’s brain were more MNase sensitive than the larger expansions. Quantifying MNase inaccessibility between asymptomatic and symptomatic brains is hampered by the repeat length variations they presented, which we have shown to affect chromatin accessibility (Fig. 5D). Expression of the *C9orf72* transcript in the asymptomatic *C9orf72*Exp father was dramatically increased 2.5-7-fold in the frontal cortex, cerebellum, spinal cord, blood, and four other tissues compared to the affected daughter’s tissues, whose *C9orf72* levels were twice less than an individual devoid of expansions (29, 88). We suggest that the increased transcription in the father’s brain arose from the shorter range of expansion sizes (13-1200 repeats) which were less MNase resistant than the longer expansions, which is consistent with both the over expression in the father’s blood, which contained predominantly 70 repeats, and with the low transcription in the daughter’s brain that contained only very large expansions. This interpretation is consistent with the 2.5-5-fold increased expression of intermediate *C9orf72* repeats in human brains, blood, and knockin iPSC-derived neural progenitor cells (66, 90). This parallels the 5- to 10-fold increased expression from the premutation CGG expansions at the *FMR1* repeat in hemizygous males, which also show compact chromatin (91, 92). We conclude that *C9orf72* expansions and flanks assume a highly compact chromatin in cells and brain regions of *C9orf72*Exp carriers. MNase resistance is evident on *all C9orf72*Exp alleles in a given sample, which differs from the 7-14% of *C9orf72*Exp cells cytogenetically showing FRA9A-Chr9s, like FXS, only a portion of the CGG-expanded ChrXs show FRAXA (69, 70). In this manner, MNase resistance might be considered a proxy for fragility. Towards assessing whether MNase resistance was linked to the unusual chromatin compaction of cytogenetic fragility or merely linked to expanded repeats, we assessed the

MNase accessibility of the *HTT* locus in Huntington’s disease patient cells with (CAG)21/180, which is known to not be a cytogenetically detectable fragile site (93). MNase treatment and Southern blotting resolved the mutant and non-mutant *HTT* alleles and both showed the same MNase accessibility when most of the genome was sensitive (Fig. S5, lanes 3-5), suggesting that MNase inaccessibility does not extend to this non-fragile mutant locus. As part of another study, we extended the MNase inaccessibility to the CGG-expanded *FMR1* gene the molecular cause for the fragile site FRAXA. We find MNase resistance of only the CGG expanded allele (in cells from male and female carriers of the CGG expansion) (not shown). That the CGG expanded *FMR1*, which forms FRAXA, but not the non-fragile CAG expanded *HTT*, forms an MNase resitant region supports our suggestion that MNase inaccessibility of the *C9orf72*Exp can be considered a proxy for chromosomal fragility. We suggest that the MNase resistance of the *C9orf72*Exp locus in human brains is a reflection of chromosomal fragility (Fig. 5F).

### Increased Chr9-containing micronuclei in *C9orf72*Exp cells

Micronuclei (MN) are indicators of chromosomal double-strand breaks (DSBs) including fragile sites (94–97), where MN often harbor broken chromosomes to be eliminated or reintegrated. Nuclear buds (NBuds) are chromatin-containing protrusions linked to the nucleus by a nucleoplasmic stalk (95, 97). Both MN and NBuds can arise from chromosome breaks. NBuds can become MN (95). These subnuclear structures can be enriched with tandem gene amplifications and satellite repeats (67, 98, 99). Cytoplasmic release of MN DNA triggers an immune response via DNA sensing by cGAS-STING that activates *IFN* expression (94), a pathway hyperactivated in vulnerable neurons of *C9orf72*Exp carriers (4, 12–14). Given these connections, we assessed MN and NBuds and whether they contained Chr9 in *C9orf72*Exp cells. *C9orf72*Exp cells showed elevated MN/NBud levels compared to control cells, which were further increased with folate stress (Fig. 6A). More NBuds over MNs contained Chr9 in *C9orf72*Exp cells compared to control cells. The modest increase in MN with Chr9 could be due to loss of fragile chromosome-containing MN during folate- perturbation, as suggested for FRAXA cells (Fig. 6B) (100, 101). Depending upon whether the breakpoint is centromeric or telomeric of *C9orf72*, the presence of the *C9orf72*- containing FISH signal alone or the 9q-arm FISH signal alone, supports subnuclear inclusion of an acentric or truncated Chr9, respectively, while the presence of both *C9orf72*- and q-arm signals supports the inclusion of truncated Chr9 or the full-Chr9 (Fig. 6B). An absence of either FISH signal could reflect an absence of any Chr9, or an acentric Chr9 broken telomeric of *C9orf72*. When Chr9 signals were present in MN/NBuds, the proportion of the *C9orf72* signal alone, 9q-arm alone, or the two together occurred at 1:1:5, respectively. Increased Chr9-containing MN/NBuds were evident in *C9orf72*Exp cells of presymptomatic and symptomatic individuals. Cumulatively, these data support increased chromosomal damage in *C9orf72*Exp cells.

**Figure 6.**
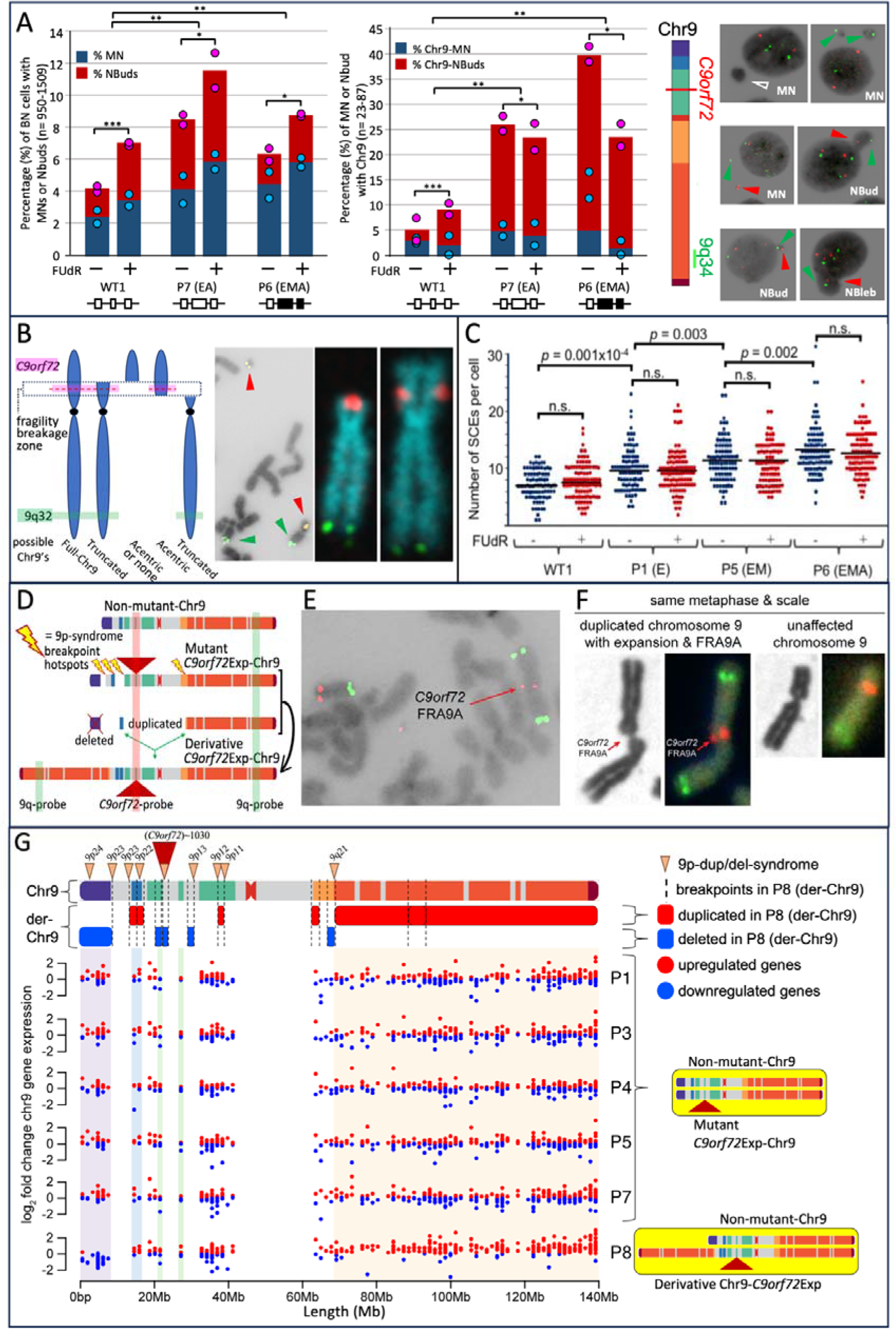
C***9***orf72Exp **cells display chromosomal instability.** (A) Quantification of micronuclei (MN) and nuclear buds (NBud) in binucleated cells without FISH, left, and with *C9orf72* and 9q-FISH probes, middle, in WT and *C9orf72*Exp cells (P6 (ssc) and P7) containing similar sized repeats (∼770) but differing methylation status. Data are means of two independent experiments. Dots show data of each experiment. Statistical analyses using Fisher’s exact test (*P<0.05, **P<0.01, ***P<0.001). Representative images of MN and NBuds in cytokinesis-blocked cells treated with FUdR for 24 hours, that contain no FISH signal (hollow arrows), a *C9orf72*-signal (red arrows), or 9q-signal (green arrows). (B) Depending upon site of breakage, *C9orf72*-FISH probes can have varying interpretations, left. Example of *bona fide* chromosome breaks in *C9orf72*Exp line P4 with acentric terminal *C9orf72*-containing fragment in same metaphase spread as a truncated-Chr9 that is free of *C9orf72*-signal, middle. Example of truncated-Chr9 with *C9orf72*-signal, and control Chr9 from same spread, right. (C) Levels of global sister chromatid exchanges (SCEs) in WT and *C9orf72*Exp cells +/-FUdR. Methylation/clinical status is as per Figure 3. Each datapoint represents a single metaphase. At least 100 mitotic cells were analyzed per condition in each experiment. Statistical analyses were performed by student’s t-test (P<0.01). NS: not significant. See also Figure S5B-E. (D) *C9orf7*Exp line P8 with complex highly rearranged derivative-Chr9, schematic, see text and Figure S7A. (E) Example of FRA9A-expressing derivative- Chr9 and control Chr9 in same spread with *C9orf72*- (red) and 9q-FISH probes (green) (100%=100/100 metaphases). (F) Only the derivative-Chr9 expresses FRA9A coincident with the *C9orf72*-FISH signal, revealing derivative-Chr9 to be the *C9orf72*Exp-Chr9, (+FUdR, 7%=7/100, 99% confidence). (G) Breakpoint analysis (top) and gene expression (binned/gene) along Chr9 in each of the *C9orf72* Exp cells with trisomic (red dots) and monosomic (blue dots) in duplicated and deleted regions of the derivative-Chr9 (P8). Breakpoint and copy number change analysis in P8 was by whole genome sequencing, see Table S3. An interactive html file detailing mis-expressed genes can be accessed here: https://data.cyverse.org/dav-anon/iplant/home/ljsznajder/FRA9A/Chromosome9.html), also presented in Table S4. See Figure S7B-D for statistical significance of differential gene expression.

We also observed increased nuclear blebs (NBlebs) in *C9orf72*Exp compared to control cells (Fig. 6A). NBlebs are chromatin-containing nuclear herniations (95, 97, 102, 103), without an obvious constriction between the nucleus and the protruding nuclear material. NBlebs may in some instances lead to NBuds, which may lead to MN (95). NBlebs are hallmarks of the DNA damage accumulating progeroid syndromes and can be enriched with genomic regions with perturbed chromatin and DNA damage (γH2AX) (102–106). In *C9orf72*Exp cells, NBlebs contained Chr9 at higher levels than in cells without expansions (Fig. S6A), which is consistent with reports of NBlebs being enriched with regions having altered chromatin compaction and damaged DNA (102–104).

### *C9orf72*Exp and double-strand breakage

Direct support that fragile sites are prone to DNA breakage is the cytogenetic manifestation of a truncated chromosome or acentric chromosome fragment broken at the fragile site (107, 108). We observed truncated Chr9s broken at 9p21, where the truncated p21→pter acentric arm, containing the *C9orf72*-FISH signal, remained within the *same* metaphase spread (Fig. 6B). We observed numerous instances of truncated Chr9s with the *C9orf72* signal at the broken end, in the absence of their acentric p21→pter acentric arms, which had likely been lost. 9p21-truncated Chr9s without *C9orf72*-signal were also observed, consistent with FRA9A’s broad ∼8.2Mb breakage zone (Fig. 6B, see truncated Chr9 broken centromeric and telomeric of the *C9orf72*-FISH signal). Our cytogenetic findings, coupled with the increased Chr9-containing MN/NBuds, is convincing evidence that double-strand DNA breaks occur at FRA9A, consistent with the γ-H2AX foci in cells, spinal cord, and brains of *C9orf72*Exp carriers (23, 24).

We further explored the susceptibility of GGGGCC repeats to double-strand DNA breaks using a yeast system that quantifies rates of chromosome end-loss (109). A (GGGGCC)60 inserted into a yeast chromosome was extremely unstable, yielding mixed populations with 15-60 repeat units (Fig. S7A-D). Rates of breakage showed a significant length-dependent increase: Tracts of 35-60 repeats broke 6-16-fold compared to (GGGGCC)15, which incurred breaks at 10-fold higher rates than a (CAG)85 tract analyzed in the same assay (Fig. S7B and Fig. S7C). Thus, even short GGGGCC repeats can be highly prone to DNA breakage. Instability of a (GGGGCC)70 tract in humans was reported (29).

### Sister-chromatid exchange and recombinant events in *C9orf72* expansion cells

An increased incidence of sister-chromatid exchanges (SCEs) occurs at rare and common fragile sites (110). We analyzed SCE formation in *C9orf72*Exp. Unexpectedly, *C9orf72*Exp cells exhibit an increase, up to 2-fold, in spontaneous genome-wide SCEs/cell (∼7 up to 10- 15/metaphase), compared to control cells without expansions (Fig. 6C). Increased SCEs were evident in *C9orf72*Exp cells from presymptomatic and symptomatic individuals (Fig. 6C, compare P1, P5 with P6, all greater than WT). The increased SCEs did not require FUdR. This increase is significant, yet modest relative to 10-fold increased SCE/metaphase in individuals with the chromosome instability Bloom syndrome. Bloom individuals present high SCEs (111–114), increased MN (112, 115), a self-DNA cGAS-STING-mediated *IFN* over-expression (115), and often succumb to cancer before age 30 due to a coincident 10-fold increased mutation rate (114, 116, 117). Chemical and genetic factors can drive SCEs (114, 118, 119). In all instances, SCEs predominantly occur at common (120–122) and rare fragile sites (110, 123–126), regardless of whether the fragile site is being induced or not (122, 127). Consistent with this, the SCEs levels in *C9orf72*Exp cells were mildly but not significantly increased by FUdR (Fig. 6C). Through FISH mapping SCEs were significantly enriched on Chr9p, where *C9orf72* resides (Fig. S6B-D). Thus, the GGGGCC-expanded FRA9A, like the CGG-expanded FRAXA, the AT-repeat-expanded FRA16B, and the EBNA1-motif repeat 11q23 fragile site, incurs increased SCEs (85, 87, 110, 126, 128). These findings extend the forms of genetic instability and the sources of DNA damage in *C9orf72*Exp cells to include global and focal SCEs.

Unexpectedly, one of the *C9orf72*Exp cell lines (P8) showed complex rearrangements restricted to one Chr9, clearly evident in every metaphase (100/100 observed) (Fig. 6D-F). Specifically, one Chr9 showed a duplication of nearly the entire 9q-arm (9qter-q21.11 fused to 9p23, Fig. 6C), a terminal deletion of 9pter-p23, and duplications of 9p23, 9p12, 9p11.2, 9q13, 9q21.11, and deletions in 9q13, making a derivative-Chr9 (see CNV and whole-genome sequencing analysis, Fig. 6D-E, Fig. S8A, Table S3). Breakpoint pattern was reminiscent to 9p deletion/duplication syndrome (Fig. 6G top) (55, 129). Importantly, only the derivative-Chr9 expressed FRA9A with FUdR in 7% (7/100) metaphases (Fig. 6F), indicating it arose from the *C9orf72*Exp Chr9. Strikingly every metaphase showed the same rearrangements, which were evident prior to FUdR treatment. In our previous examinations of thousands of metaphases from control cells, we have not observed such homogeneity of rearrangements, making it highly unlikely that this derivative-Chr9 arose by cell line establishment or culturing. It is also highly unlikely that this individual, who was neurotypical until presenting with bulbar onset ALS at age 56, had inherited this complex set of rearrangements constitutionally in all tissues, as congenitally such rearrangements would be incompatible with neurotypical life (55, 129). Rather than inherited, it is likely the rearrangements arose somatically in the blood of the *C9orf72*Exp individual, which is supported by our observation of similar rearrangements of the *C9orf72* transgene-containing Chr6 in mouse tissues (below). Whole genome sequencing revealed the derivative- *C9orf72*Exp-Chr9 presented a unique state with ≥13 breakpoints clustered on 9p, with direct and inverted junctions, confined to a single chromosome, as such, fulfilling the criteria of chromothripsis (130–132). It is possible the FRA9A-expressing *C9orf72*Exp-Chr9 isolated in a micronucleus, was shattered and reassembled as the derivative-Chr9 (133). A single experimentally targeted double-strand break can induce chromothripsis to the targeted chromosome (134, 135). The derivative-Chr9 may have arisen by stepwise alterations. The origin of the derivative-*C9orf72*Exp-Chr9 could not be discerned as the patient was deceased. The complex chromosomal rearrangements of the mutant *C9orf72*Exp FRA9A-expressing Chr9 could have downstream impact.

While we cannot exclude the possibility that the highly rearranged derrivarive-Chr9 in the *C9orf72*Exp cell line (P8) arose during cell line establishment or passage, it is difficult to argue away the coincidences that the rearrangements we observe were limited to only one Chr9 – the Chr9 with the *C9orf72*Exp expressing FRA9A (Fig. 6D-F), and that the breakpoints were non-random but aligned with the breakpoints of 9p-duplication/deletion syndrome (Fig. 6G) (55). We also observed numerous forms of chromosome rearrangements of murine chromosomes harboring the human *C9orf72* expansion, including one reminiscent of the der-Chr9 we observed in the human cell (see murine cytogenetics described below). Moreover, the cytogenetic observation of thousands of human EBV-transformed cells have not presented such chromosomal instability (59), asides from the EBV-induced fragility at 11q23, which depends upon the EBV protein EBNA1 (84–87).

Gene expression across most of the derivative-Chr9 is dysregulated. Expression analysis by RNA-seq comparing P8 to other *C9orf72*Exp cells, revealed trisomic and monosomic expression levels of many genes corresponding to the large duplicated and deleted regions (Fig. 6G, see red and blue dots, for significance see Fig. S8B-D, an interactive html file detailing mis-expressed genes can be accessed here: https://data.cyverse.org/dav-anon/iplant/home/ljsznajder/FRA9A/Chromosome9.html), also presented in Table S4. Amongst the dysregulated genes along the der-Chr9 in P8, many are also dysregulated in vulnerable neurons of *C9orf72*Exp ALS patients, including ALS- and inflammation-linked genes (https://www.biorxiv.org/content/10.1101/2023.12.22.573083v1, Table S4). Breakpoint and copy number changes analyzed in P8 are presented in Table S4.

### Cell culture in FUdR increases repeat instability

We tested whether repeat length instability could be generated in *C9orf72*Exp patient cells under fragile site inducing conditions. We observed that expansion tracts shifted towards *shorter* repeat lengths over culture time in FUdR-treated cells (Fig. S9). In contrast, under uninduced conditions, the expanded repeat showed a general shift towards *longer* repeat sizes or the more predominant repeat length (Fig. S9). FUdR affected the length of only the mutant *C9orf72*Exp allele. FUdR-induced changes were consistent in three different *C9orf72*Exp cell lines and did not appear to depend upon methylation or clinical status (Fig. S9). These results support repeat size variations being generated under folate-perturbing fragile site inducing conditions.

### Tissue-specific somatic repeat expansions, chromosomal fragility, and chromosomal instability in *C9orf72*Exp mice

Tissue-specific somatic repeat instability, FRA9A chromosome fragility, and chromosome instability, was recapitulated in *C9orf72*Exp-Transgenic(Tg) mice (*C9orf72*Exp-Tg) harboring a single copy of the human *C9orf72* with a GGGGCC expansion integrated into murine Chr6 (136). Using this mouse as a tool for instability, we isolated cells from various organs of *C9orf72*Exp-Tg and control non-Tg mice, aged 5-months, when cell proliferation in the CNS and peripheral organs had ceased at 2-months of age (137–139) (see Methods). To retain as close a representation of cell types, harvested tissues were cultured for short periods sufficient for chromosome preparation (1-4 weeks, with no storage). Tissues from three mice inheriting ∼620-690 repeats (tail at two-weeks, aged 5-months) showed a heterogenous smear of 586-1488 repeat sizes, with some of the greatest expansions in the CNS (Fig. 7A). This supports massive post-natal inter- and intra-tissue repeat expansions in non-proliferating murine tissues, as observed in *C9orf72*Exp humans (26–30). Expansions were evident as early as 2-weeks (Shum *et al.*, in preparation).

**Figure 7.**
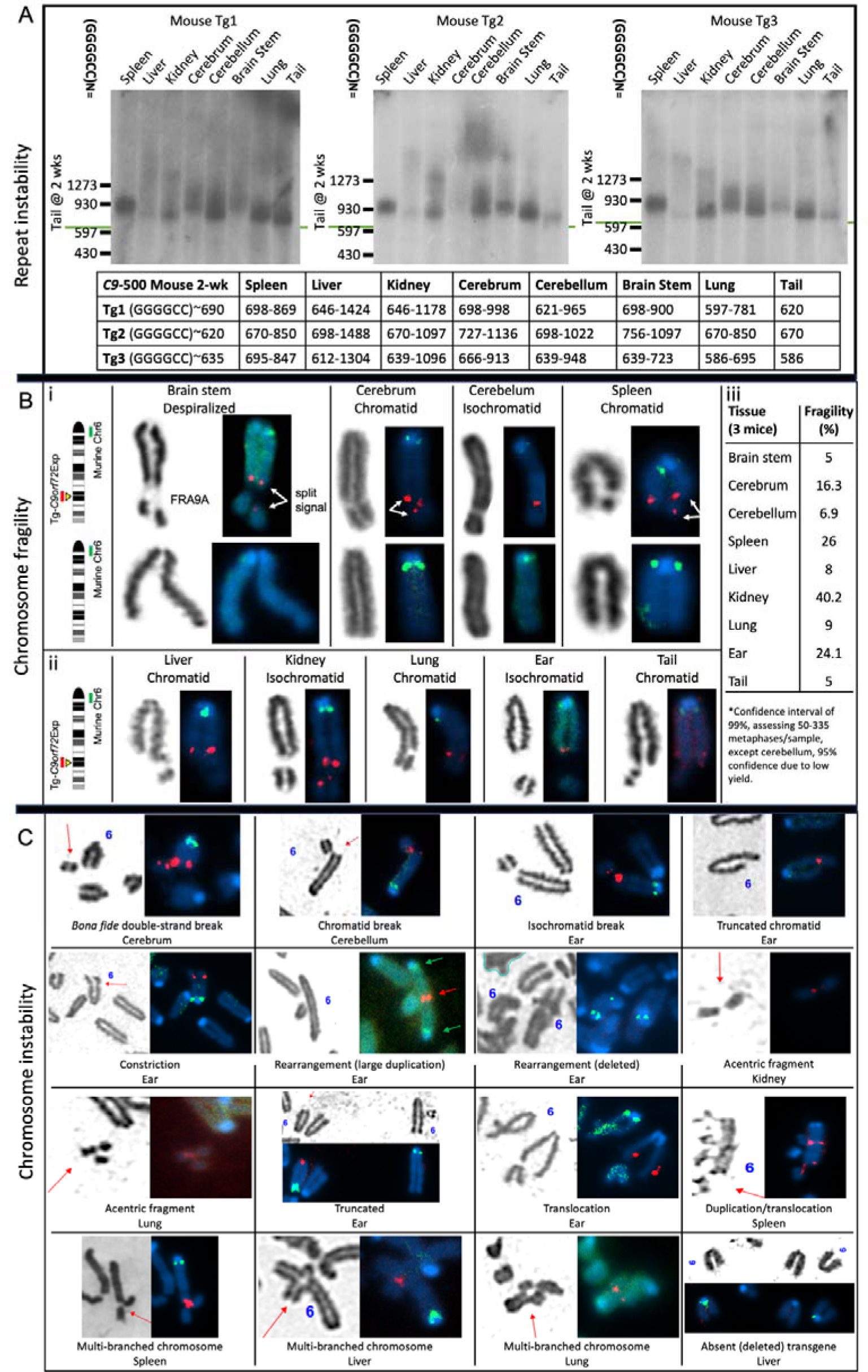
C***9***orf72 **repeat instability, chromosomal fragility and chromosomal instability in *C9orf72*Exp-Tg-mice** Southern blots for *C9orf72* transgene in C9-500 mouse tissues. C9-500 mouse tissues were sized for repeat size via Southern blot in three *C9orf72*Exp Tg-mice. The tail was the most stable tissue. (A) Somatic repeat instability by Southern blots in tissues of three 5-month-old *C9orf72*Exp-Transgenic mice harboring a single copy of the human *C9orf72* gene with a GGGGCC expansion integrated into murine Chr6 (136). (B) Cells isolated from tissues of the mice shown in panel A were grown for 1-4 weeks sufficient for metaphase spreads for fragility analysis with 1µM FUdR with FISH analysis. A murine-specific green probe identified mouse Chr6, and human-specific red probe identified the *C9orf72*Exp transgene. Pictured are FSFS in the CNS (i), the periphery (II), and quantified in panel iii. (C) Chromosomal instability at the *C9orf72*Exp transgene was assessed in the same tissues used in panel C, using the same FISH probes.

Chromosomal fragility of the *C9orf72*Exp transgene occurred in all tissues at frequencies of 5% to >35% (99% confidence, Fig. 7Bi-iii). Fragility presented as despiralized regions, constrictions, chromatid, and isochromatid, breaks, with split transgene-specific *C9orf72* FISH signals, as we observed in human *C9orf72*Exp cells (Fig. 5Bi). Only chromosomes from the transgenic mice showed FUdR-induced fragile sites, where fragility localized to the expanded repeat of the integrated transgene, but never to the non-transgenic Chr6 (Fig. S10). Thus, FRA9A fragility can be transferred to this Tg-mouse thereby validating the *C9orf72* repeat expansion as the molecular cause of FRA9A.

Chromosomal instability was evident on the *C9or72*Exp-Tg-containing Chr6, including acentrics, truncations, duplications, translocations, deletions, and multi-branched chromosomes (Fig. 7C). In one instance the *C9orf72*Exp-Tg translocated itself from murine Chr6 to another chromosome (Fig. 7C), and in another the *C9orf72*Exp-Tg was deleted (Fig. 7C). We observed a rearranged chromosome with a large duplication (Fig. 7C), reminiscent of the derivative-*C9orf72*Exp-Chr9 in ALS patient, P8 (Fig. 6D-G). Chromosomal fragility and instability were observed in all tissues of all three *C9or72*Exp-Tg mice.

While rearranged chromosomes were rare relative to fragility, neither were ever observed in the non-transgenic murine Chr6 at 6qE3, the site of transgene integration, or in any control non-transgenic tissues (three replicate mice). The absence of fragility at 6qE3 in control non-transgenic tissues is consistent with 6qE3 not being an endogenous fragile site in mice (140).

Together our data support the ability of the *C9orf72*Exp-Transgene to incur post-natal somatic tissue-specific repeat expansions, display chromosomal fragility and chromosomal instability, as in humans. Our findings provide insight into fragile site expression and rearrangement in the CNS and periphery.

## DISCUSSION

Here we demonstrate that *C9orf72*Exp is the molecular cause of the rare folate-sensitive FRA9A, cytogenetically observed more than 4 decades ago (58, 141). This reveals previously unconsidered avenues to study genotype-phenotype relationships for co-occurring disease states associated with *C9orf72*Exp and attributes linked to the 9p21 region. First, folate- sensitivity suggests a possible gene-environment interaction. Second, FRA9A may serve as a DNA source to activate DDR and *IFN* pathways. Third, FRA9A chromosomal instability and global SCEs may increase somatic mutations at 9p21 and beyond.

### Disease penetrance & gene-environment

Gene-environment interactions have long been suspected for ALS, FTD, and the incompletely penetrant co-occurring phenotypes of *C9orf72*Exp (142). Folate-sensitive induction of *C9orf72*Exp repeat instability, brain region-specific repeat expansions, FRA9A fragility, chromosomal instability, and methylation-sensitive chromatin compaction, which we have shown can arise in cells and tissues of both presymptomatic and symptomatic individuals, may have clinical implications. Lifestyle, including nutritional choices, of *C9orf72*Exp carriers can affect disease onset and progression (40). Proper levels of the linked vitamins B9 (folate) and B12 are protectors from ALS (143). Individuals with folate deficiencies, nutritionally-, pharmaceutically-, or genetically-induced, are associated with chromosomal fragility, localized chromosomal rearrangements (144–153), megaloblastic/pernicious anemia, ALS- and FTD-like symptoms (154, 155), reviewed (59). In such cases these presentations were reversed upon B12 supplementation – indicating that chromosomal instability was induced by folate-stress. Somatic GGGGCC repeat instability, FRA9A fragility, and global increase of SCEs depend upon *C9orf72* expansion and are folate-sensitive and can occur in cells from presymptomatic *C9orf72*Exp individuals.

*C9orf72*Exp carriers, in folate-deficient states, may be predisposed to repeat instability, FRA9A fragility, and chromosomal instability, which if exacerbated over time may impact gene dysregulation, altering the susceptibility of phenotypic variations amongst *C9orf72*Exp carriers. Fragile sites and SCEs can be induced in the blood and myeloid cells by lifestyle choices, including cigarette smoking, alcohol consumption, and diet (59, 156–160). Tobacco and alcohol consumption by presymptomatic *C9orf72*Exp carriers affected disease age-of-onset (40). Various clinically used agents (161), including some considered for use in ALS/FTD, *C9orf72*Exp (162), and fragile X syndrome (163), like phenytoin or decitabine can induce fragile sites (59, 164), induce sister-chromatid exchanges (165, 166), and modify chromosome condensation (167, 168). Age-dependent ongoing repeat expansions in the brain have been suggested as a contributor to disease (26, 88, 169, 170), could be folate sensitive. Additionally, CpG-methylation of *C9orf72*Exp (169, 171) and acceleration of genome-specific DNA methylation age (epigenetic clock) are modifiers of disease onset and progression in *C9orf72*Exp carriers (172, 173). Chromatin compaction of *C9orf72*Exp is sensitive to the degree of expansion and CpG-methylation (Fig. 5). Efforts to prophylactically slow the epigenetic clock are a central focus (174–176). The *C9orf72* expansion was recently reported to be bound by the chromatin remodeler DAXX (177). Coincidentally, DAXX, is known to associate with satellite and telomere repeats and its downregulation led to a global increase in MNase sensitivity (178–180). Environmental contributions to *C9orf72*Exp diseases must be considered.

Various genetic, molecular, and environmental aspects have been demonstrated to modify disease presentation in *C9orf72*Exp carriers (3, 4, 7, 8, 40, 42, 45, 181–188). The 90-year old male *C9orf72*Exp carrier presented herein (Fig. 5) is likely to be truly asymptomatic, as a long presymptomatic phase with evident degeneration often detected decades before ALS or

FTD symptom manifestation (238, 239), pathology that was not present in his brain (88). Clinical differences between the truly asymptomatic father and symptomatic daughter studied herein could not be explained by either genetic modifiers (*TMEM106B*, *ATXN2*, Finnish haplotype), both expressed similar burdens of p62 inclusions, similar burdens of sense and antisense RNA foci, similar burdens of all five types of RAN-peptides (GA > GP > GR > PA/PR, except for the cerebellum, which showed more RAN-peptide inclusions (∼7-fold) and sense RNA foci (∼2-fold) in the asymptomatic father compared to the ALS/FTD-affected daughter) (29, 88) (summarized in Table S2). Also, neither lifestyle choices, life events, nor autoimmunity, could explain their phenotypic differences (29, 88) (Table S2). However, compared to the daughter, in the brain the father showed extreme repeat length mosaicism (13-3500 repeats) (Fig. 5D), with increased MNase accessibility of shorter expansions, increased *C9orf72* expression (mRNA and C9ORF72-long, but not C9ORF72-short protein) (29, 88), increased TDP-43 inclusions (29, 88), and an epigenetic age ∼9 years younger than his chronologic age (the daughter was 3 years older) (88, 172) (summarized in Table S2). Thus, these somatic variations deserve further attention in pre-, asymptomatic, and symptomatic *C9orf72*Exp carriers.

### FRA9A as a DNA source to activate the DDR and *IFN* pathways

Both a spontaneous activated DNA damage response (DDR) (15–25) and a hyperactivated *IFN* cGAS-STING-associated immune response occur in *C9orf72*Exp patient cells and brains (4, 12–14). The source of the damaged DNA or DNA-immunostimulant to activate either DDR or *IFN* response is unknown. DDR in cells and brains of *C9orf72*Exp, evident as increased γ-H2AX, phosphorylated-ATM, cleaved PARP-1, and 53BP1, is coincident with DSBs (22–24). We reveal that DSBs arise at *C9orf72*Exp via heightened FRA9A fragility, broken/rearranged Chr9s, increased genome-wide SCEs, increased MN, and Chr9-containing

MN. The increased C9ORF72 protein levels in multiple tissues of the asymptomatic father shown to have massive mosaic somatic expansions (Fig. 5F, Table S2 and (29, 88)) may suppress an autoimmune response by increasing STING levels. Models of the *C9orf72*Exp- linked DDR activation involve RAN-peptide induced p53-target gene activation, inhibition, or mislocalization of the DSB DNA repair proteins ATM, 53BP1, TDP-43, FUS, Ku80, and PARP-1 (15–25). Other non-RAN-peptide DDR paths have received little attention.

The unusual >33kb of compact chromatin at FRA9A’s core fragility zone (Fig. 5A), could be susceptible to DSBs, which would contribute to DDR activation. Supporting this possibility is that experimentally-induced locus-specific compacted chromatin of an artificial ∼10kb stretch of ∼250 tandem arrays of LacO, which becomes a fragile site, has been demonstrated to activate a DDR in the absence of damage (189–192). A contribution of FRA9A fragility to DDR can act in parallel to RAN-peptide induced DDR (23, 24). Disruption of higher-order chromosomal organization may also contribute to DDR and *IFN* responses. Long-range chromosomal interactions involving 9p21 are activated in the inflammatory response. All *IFN* genes at 9p21 are transcriptionally inactive until immunostimulated by viral infection or cytosolic self-DNAs, whereupon dramatic chromatin (193) and multiple intra-9p21, inter-arm (9p21-9q34.2), and inter-chromosomal (9p21-4p14 and 9p21-18q21.1) interactions arise to activate *IFN* cluster expression and inflammation (47, 194). Reduced levels of the C9ORF72 protein in *C9orf72*Exp patients, being unable to supress STING, leads to a sustained hyperactivation of *IFN* in brains (195). Upon FRA9A breakage the *C9orf72*Exp-Chr9 activates DDR and is then sequestered to MN whose contents can be released to the cytosol to activate *IFN* through cGAS-STING, and this sequestration may disrupt other inter- and intra-Chr9 interactions (196). That disruption of intra-/inter-chromosomal associations can drive DDR and *IFN* cascades is supported by the experimental disruption of such interactions leading to chromosome sequestration to MN (197, 198). CGG expansions at *FMR1*, the cause of FRAXA, disrupt intra- and inter-chromosome interactions (199). The 2-fold increase in global SCEs in *C9orf72*Exp cells we observe may contribute to DDR and MN. MN enriched with the DSB repair protein, TDP-43, have been detected in brains of ALS patients (200). Our suggestion that DNA damage induced by FRA9A fragility can serve as a DNA-immunostimulant in the CNS is consistent with reports of *in vivo* DNA damage by ATM-deficiency or brain trauma that activates an *IFN* response in motor neurons (201–203). Our working model that the *C9orf72*Exp acts as a chromatin compacted FSFS and is an inducer of DNA damage at many locations including on Chr9, is suited to non-cycling neurons.

We suggest that the cytoplasmic release of DNA from FRA9A-enhanced MN, may further tip the balance by exacerbating the stimulation of the aberrantly hyperactive cGAS-STING initiated *IFN* expression in *C9orf72*Exp carriers (4, 13, 14) – which may explain the co- morbid partially-penetrant autoimmunity (1–6). FRA9A broken-Chr9s and increased global SCEs might also further exacerbate the DDR cascade, as DNA-immunostimulation enhances sense and antisense transcription from *C9orf72* (42) whose RAN-peptide products would feedback upon the DDR by suppressing DSB DNA repair proteins ATM, p53, TDP-43, FUS, Ku80, and PARP-1 (15–25). The neuroinflammatory and autoimmune phenotypes of *C9orf72*Exp carriers is complex (14), and the effects of either FRA9A altered chromatin, or MNs containing broken Chr9s upon the *IFN* response is unknown.

### FRA9A chromosomal instability, mosaic somatic mutations, and 9p21-associated diseases

A reasonable prediction from our findings is that the *C9orf72*/FRA9A fragile site may, like other fragile sites, predispose to mosaic deletions, duplications, or rearrangements resulting in altered expression of genes on 9p21 or throughout Chr9. Such mosaic chromosomal variations may vary between tissues, accumulate with age, and lead to phenotypic variability. In the case of *FMR1*/FRAXA, phenotypic variability can arise through mosaic genetic or chromosomal instability (deletions, duplications, and rearrangements) extending beyond the *FMR1* CGG-repeat, as revealed through decades of isolated case reports (59). Similar phenomena may arise at the mutant *C9orf72*, as suggested by our observation of truncated- Chr9s broken at *C9orf72*, acentric-Chr9s, chromosomal instability of the *C9orf72*Exp allele, and heightened levels of genome-wide SCEs. Detection of somatic mosaic levels of chromosomal instability will likely require more focused studies. For example, the dramatic Chr9 rearrangements, evident by FISH in every metaphase of the ALS/*C9orf72*Exp line P8, escaped detection by repeat-primed PCR and by Southern blot assessment of the repeat size (compare Fig. 3B with Fig. 6F). Indeed, mosaic mutations at various loci (reviewed in (204–206)) including *FMR1* (207–212) required focused attention. Mosaic mutations may be confined to specific cell types, as recently revealed for Purkinje cells with large somatic CAG-expansions, amongst stable tracts in the cerebellum of Huntington’s disease patients (213, 214). One might expect an increase in somatic mutation rates throughout the genome in *C9orf72*Exp patients, as the increased 2-fold global SCEs we observed may have a coincident increase in mutation rate, similar to the 10-fold increased mutation rates in BLM individuals who show a 10-fold increase of SCEs (111, 112, 116, 117). *C9orf72*Exp cells from presymptomatic individuals show folate-sensitive repeat instability, FRA9A fragility, chromosomal instability, MNase-resistance, increased levels of micronuclei/NBuds, and global sister chromatid exchanges. Long-term display of these features may exacerbate disease-onset and affect penetrance. Our findings support that the *C9orf72*Exp patients may incur mosaic mutations at under-appreciated levels, and recent findings support increased somatic mutations in brains of *C9orf72*Exp individuals (https://www.biorxiv.org/content/10.1101/2023.11.30.569436v1).

Knowledge gaps remain for FRA9A/*C9orf72*. In particular, the co-occurrence of many diverse phenotypes in *C9orf72* expansion carriers is unclear. That FRA9A encompasses the *IFN* cluster, *CDKN2A*, *IFNk*, *LINGO2*, *TOPORS*, *APTX*, *DNAJA1*, and *SMU1* genes, each independently associated with symptoms that can co-occur in *C9orf72*Exp carriers, provide new insights into the possible involvement of chromosomal instability (Fig. 1). An increase in ALS occurring in cancer survivors, particularly melanoma, has been repeatedly reported (7, 215–219). While a correlation of *C9orf72*Exp in cancer predisposition has not been studied in large populations, *C9orf72*Exp carriers have a significantly higher risk of melanoma (7). *C9orf72*Exp carriers may succumb to cancer prior to onset or awareness of ALS. That both *CDKN2A* and the *IFN* gene cluster, frequently deleted in melanoma, are encompassed in FRA9A’s fragility zone introduces the possible involvement of *C9orf72*Exp. Mosaic somatic mutations that extend beyond the *C9orf72* repeat, may affect expression of genes encompassed by FRA9A or beyond. Thus, revisiting the *C9orf72*/9p21 locus with the awareness that it is, and can share the attributes of, a chromosomal fragile site should provide key insights into the genotype-phenotype relationships for the many and varied presentations associated with the 9p21 region.

## MATERIAL AND METHODS

### *In silico* datasets and analyses

Sequence feature and epigenetic datasets used in Figure 2 and Figure S1 are described as follows. Repeat annotations and %GC tracks were obtained from the UCSC genome browser (hg19). Wavelet-summarized repliseq data (ENCODE) was obtained in bigWig format from GEO:GSE34399. Erythroblast SNS-seq data and replication origins were obtained in bedGraph and bed format, respectively, from GEO:GSE6197. Imputed %DNA methylation for GM12878 was obtained from the roadmap epigenomics data portal: (https://egg2.wustl.edu/roadmap/data/byFileType/signal/consolidatedImputed/DNAMethylSBS/E116-DNAMethylSBS.imputed.FractionalMethylation.signal.bigwig). Pre-calculated 2nd-order DNA shape features were obtained from the Rohs lab GBShape database.

### Cell culture and fragile site induction

All cytogenetic studies were performed using Epstein-Barr virus transformed lymphoblastoid cell lines from *C9orf72-*ALS patients, expansion carriers, and control individuals provided by Dr. Guy Rouleau (McGill University). One of the patients, P4, displayed signs of FTD. All patients provided written informed consent. Cells were cultured in RPMI 1640 media supplemented with 15% FBS (ThermoFisher Scientific), 2 mM L-glutamine, Pen/Strep (100 IU/mL, 100 µg/mL), and 1 mM sodium pyruvate. Two modes of FSFS induction were used to demonstrate fragility at 9p21.2: i) 24 hours with 0.1 µM 5-fluorodeoxyuridine (FUdR) or ii) 14 days with Media 199 (Gibco, Cat #: 11150-059) supplemented with 2% FCS, 2 mM L-glutamine, and Pen/Strep (100 IU/mL, 100 µg/mL). Cells in 199 media were treated with 100 nM methotrexate for 16 hours, washed and then treated with 0.01 µM bromodeoxyuridine (BrdU) for a total of 3 hours prior to harvest. All cells were treated with 6 µg/mL ethidium bromide (EtBr) and 0.05 µg/mL KaryoMAX (Gibco; Cat #: 15212012) prior to harvest. Cells were then centrifuged and resuspended in 0.075 M KCl solution for 30 minutes at 37°C followed by several washes with fixative solution (3:1 methanol:acetic acid). Long-term FUdR culture experiments were performed in either RPMI 1640 media (as described above) or RPMI 1640 media + 0.1 µM FUdR for a total of 8 weeks. Media and FUdR was replaced every 2-3 days. Cells were harvested at 1, 2, 4, 6, 8-week time points for DNA purification and Southern blot was run as described below.

### Metaphase chromosome and FISH analysis

Slide preparation for both cytogenetic and FISH analysis of metaphase spreads were made from fixed cell suspensions using a Model CDS 5 Cytogenetic Drying chamber according to standard methods. For 199 media conditions, breaks and gaps were quantified on total Giemsa stained metaphases and subsequently G-banded to karyotype each metaphase and confirm fragile site on 9p-arm. SpectrumGreen & SpectrumOrange-conjugated dUTP probes (RP11-15P13, RP11-274I21, RP11-1149M23, RP11-491J7, RP11-945G22, RP11-1079P2, RP11-360B21, RP11-696J10) from RPCI-11 human BAC library were generated by The Centre for Applied Genomics (TCAG, PGCRL, Toronto, Canada) using the Vysis nick translation kit (Cat #07J00-001; Abbott Molecular, Illinois, USA) in accordance with manufacturer’s instructions. Chromosomes were counterstained with DAPI. One hundred metaphases per sample were scored for each condition using either brightfield microscopy, Olympus IX81 quorum spinning disk confocal microscope, or Zeiss Axioplan 2 fluorescence microscope. Images were collected using a CCD camera (Hamamatsu C9100-13 EM-CCD, Hamamatsu, Japan) (Princeton Instruments Pentamax, Roper Scientific, New Jersey, USA) and analyzed using Volocity (Perkin Elmer, Massachusetts, USA) or CytoVision (Leica Biosystems, Germany) software.

### Southern blot analysis of *C9orf72* repeat expansion size

Southern blot analysis was performed to size *C9orf72* repeat expansions as this method is the gold standard for sizing large (GGGGCC)n repeat expansions (220). Southern blotting provides an estimate of non-bias expansion sizes (221). We used Southern blotting method (220, 222) with 241 bp probe, which anneals 153 bp upstream of the (GGGGCC)n repeat tract, and used selected restriction-endonuclease digestions of genomic DNA (as noted in the figures) – which permit sensitive detection of repeat size and length heterogeneity at molar levels. The 241 bp probe was produced by PCR using primers (Fwd: 5’– AGAACAGGACAAGTTGCCCC–3’ and Rev: 5’–AACACACACCTCCTAAACCC–3’) as published (220, 222). For mapping boundaries of MNase accessibility an additional probe was used (see below). Molecular weight markers were used to estimate the size of repeats. Distances between molecular weight markers were measured using ImageLab software (BioRad) and plotted against the known number of base pairs for each marker. A best-fit linear regression was calculated in Microsoft Excel and the equation of this line was used to calculate the size of each band on the blot. Different restriction enzyme combinations were used to cut the genomic DNA, as noted in each figure, hence resulting in different flank repeat sizes.

### Methylation analysis of the *C9orf72* repeat

Methylation analyses were done in blinded experiments using the same DNA prep as for Southern blot. Genomic DNA was bisulfite converted using the EZ DNA Methylation- Lightning™ Kit (Zymo). We estimated the number of methylated CpG-sites at the CpG- island 5′ of the (GGGGCC)n repeat using bisulfite sequencing as reported previously (64). To estimate the methylation of the (GGGGCC)n repeat itself, we used a qualitative GGGGCC- methylation assay, sensitive enough to detect repeat methylation in a mixture containing ∼5% high-methylated DNA standard (60). We always observed methylation mosaicism for expanded repeat samples, revealed by presence of the amplification product of the expansion in both methylated (blue) and unmethylated (green) channels (223).

### Micronucleus and nuclear bud detection

Micronuclei and nuclear bud analyses were done as previously described with some modifications (95, 97, 224). Briefly, cells were treated with 0.5 µM FUdR for a total of 24 hours, with cytochalasin B (4 µg/mL) being added 3 hours after addition of FUdR. Cells were briefly incubated in 0.075 M KCl, fixed with Carnoy’s fixative and slides with sparsely spread cells were prepared using a Model CDS 5 Cytogenetic Drying chamber (Thermotron, Michigan, USA). SpectrumGreen (RP11-696J10) & SpectrumOrange (RP11-672M4)- conjugated dUTP probes from RPCI-11 human BAC library were generated by The Centre for Applied Genomics (TCAG, PGCRL, Toronto, Canada) using the Vysis nick translation kit (Cat #07J00-001; Abbott Molecular, Illinois, USA) in accordance with manufacturer’s instructions. Nuclei were counterstained with DAPI. At least 950 binucleated cells were scored per condition for the presence of micronuclei and nuclear buds and Chr9 signals, as previously described (97). Two blinded and independent experiments were carried out for each data asset. Statistical analyses were done using Fisher’s exact test for the combined values of micronuclei and nuclear bud events as they are believed to arise from the same phenomenon (97, 225). Images were captured using an Olympus BX61 fluorescence microscope. Images were collected using a CCD camera (Hamamatsu C9100-13 EM-CCD, Hamamatsu, Japan) (Princeton Instruments Pentamax, Roper Scientific, New Jersey, USA) and analyzed using ASI SpotScan software (Applied Spectral Imaging, California, USA).

### Sister-chromatid exchange (SCE) assay

Cells were incubated in the presence of 10 μM BrdU for two cell cycle periods and pulsed with 0.1 μg/mL of KaryoMAX (Gibco; Cat #: 15212012) for the last 2 hours before being harvested. FUdR (0.1 µM) was added for the last cell cycle. Harvested cells were treated with 0.075 M KCl for 20 min and subsequently fixed with methanol:acetic acid (3:1) for at least 30 minutes. Cells were fixed onto glass slides. Dried slides were incubated with 10 μg of Hoechst 33258 per ml in phosphate buffer (pH 6.8) for 20 minutes, followed by rinsing with MacIlvaine solution (164 mM Na_2_HPO_4_, 16 mM citric acid [pH 7.0]). Slides were irradiated with UV light for 2 hours and incubated in 2X SSC solution at 62°C for 1 h before staining with 5% Giemsa solution and subsequent microscopy. At least 100 metaphase cells were evaluated for SCE events genome-wide per genotype. For Chr9-specific SCEs, slides were destained with xylene solution followed by methanol:acetic acid (3:1) and then standard FISH procedures, as described above were followed using SpectrumOrange-dUTP conjugated RP11-696J10 probe. At least 30 of the same Giemsa-stained metaphases were relocalized using fluorescence microscopy and imaged to allow for quantification of SCEs specifically on the 9p- and 9q-arms.

### Construction of plasmids and yeast strains

An *Xba*I-*Hin*dIII fragment containing a region of the human *C9orf72* gene containing an expanded (GGGGCC)n repeat and 547 bp of flanking sequence (from pcDNA3.1(+)c9-200+ obtained from R.H. Brown; CF stock #435) was cloned into the MCS of plasmid pEP1 to create plasmids pEP1-(GGGGCC)n. pEP1 was made by cloning the *ADE2* gene from pRS402 into the *Bsa*AI and *Nsi*I sites of pHZ1; pHZ1 was made by cloning the G4T4 sequence into the *Sma*I site of pYIP5. The CAG repeat was cloned into pEP1 *Xho*I and *Xba*I sites. Repeat lengths in plasmid clones were confirmed by sequencing. Repeat lengths of the (GGGGCC)n plasmids were approximately 16, 29, and 36 for plasmids 504, 484, and 485, respectively; exact number of repeats varied with different sequencing runs. Plasmids were linearized with *Xcm*I and recombined into the *URA3* marked yeast artificial chromosome (YAC) CF1 in strain VPS105 (CFY #289) to create the new GGGGCC-*ADE2*-*URA3* and CAG-*ADE2*-*URA3* YACs (see Fig. S7A & S7D), which were confirmed for YAC structure by PCR. CAG tract lengths were confirmed by sequencing of PCR amplicons but (GGGGCC)n tract lengths could only be determined approximately by Southern blot since amplification from genomic DNA was unsuccessful, and are therefore reported to the nearest 5 bp.

### YAC fragility assay

End loss of the repeat-*ADE2*-*URA3* YACs was determined by an established protocol (109) except that 1 ml of an overnight culture inoculated with a specific colony was plated on media containing FOA and a low amount of adenine but lacking leucine (FOA-Leu low Ade) to determine the number of mutants, and a dilution was plated on yeast complete media lacking leucine (YC-Leu) for a total cell count. Rates were determined by the method of the median. Another 1 ml of the same culture was used to isolate genomic DNA for Southern blot to assess repeat length. For reasons not completely understood, but that could include reduction of transcription through the repeat tract, the insertion of the *ADE2* marker resulted in a lower rate of FOA^R^ compared to previously published values for the CAG-85 YAC without the *ADE2* gene. End loss was confirmed in a subset of colonies (usually one from each FOA-Leu plate) by PCR (retention of the sequence to the left of the G_4_T_4_ telomere seed and loss of sequence to the right of the repeat tract; Fig. S7).

### Southern blot - yeast

Yeast genomic DNA was digested with *Nhe*I and *Xba*I restriction enzymes, separated by horizontal gel electrophoresis, and blotted to a nylon membrane using a standard Southern protocol. Either the Roche Molecular Weight Marker VI or VII was run as a size marker. For the preparation of the probe, a 301 bp fragment adjacent to the (GGGGCC)n repeat was PCR amplified from DNA of plasmid #485 as template, using Primers #1402 (ALSsouthernFOR) and #1403 (ALSsouthernREV). Probes were labeled using the DIG High-Prime DNA Labeling and Detection Starter Kit I (Cat #: 11745832910; Roche). Repeat tract estimation was performed as previously described (Southern blot analysis of repeat expansion size).

### Chromatin accessibility *in cellulo* with MNase

Human lymphoblast cell lines from *C9orf72-*expansion carriers were assessed for MNase accessibility, as outlined in Figure 5B. Briefly, cell pellets were washed in ice-cold 1X PBS, re-pelleted, and permeabilized by ice-cold NP-40 lysis buffer (10 mM Tris pH 7.4, 10 mM NaCl, 3 mM MgCl_2_, 0.5% NP-40, 0.15 mM spermine, 0.5 mM spermidine) for 5 min on ice. Nuclei were collected at 200 X*g* (10 minutes, 4°C), washed in 1X MNase digestion buffer (10X stock: 100 mM Tris pH 7.4, 150 mM NaCl, 600 mM KCl, 5 mM spermidine, 1.5 mM spermine), re-pelleted, and subsequently resuspended in 1X MNase digestion buffer supplemented with 1mM CaCl_2_. Treatment followed with MNase for 5 min. at 22°C, at differing concentrations (refer to Fig. 5 and Fig. S4A – all serial dilutions). All digestion reactions were stopped with an equal volume of *Stop* solution (0.625 mg/mL proteinase K, 20 mM EDTA, 20 mM EGTA, 1% SDS diluted in 1X MNase buffer) and incubated at 37°C for a minimum of 1 hour. Reaction conditions were optimized to ensure over-digestion by MNase, which diminishes MNase cleavage preference. DNA was purified by phenol/chloroform extraction, RNase A treatment, and ethanol precipitation. A small amount of purified DNA was assessed for completeness of MNase digestion, by its conversion to mono-, di-, tri-nucleosome sized fragments as resolved by electrophoresis stained with ethidium bromide and visualized by UV imaging (Fig. 5B, left panel; S4A). DNA samples were analyzed by Southern blot as described.

### Chromatin accessibility in post-mortem tissues with MNase

We conducted MNase accessibility analysis using post-mortem brain tissues (orbito-frontal cortex and cerebellum) of an asymptomatic 90-year-old male and his 59-year-old ALS-FTD- affected daughter, previously deep-phenotyped (summarized in Table S2) (29, 88). The asymptomatic male carried 70-repeat *C9orf72* expansion in the blood and was deceased at 90-years of age. Tissue was Dounce homogenized using 1X MNase digestion buffer, 10% NP-40 was added to final concentration 0.5%, incubated 5 minutes on ice, filtered through a 40 μm strainer, and centrifuged. Nuclei pellets were washed in MNase digestion buffer and incubated with different concentrations of MNase. Remaining steps were performed as described above.

### Digestibility of *C9orf72* BAC DNA *in vitro* with MNase

A bacterial artificial chromosome (BAC) containing ∼160 (GGGGCC)•(GGCCCC) repeats, prepared in the lab of Piet de Jong, was restricted by *Nhe*I and *Xho*I giving rise to a shorter fragment of approximately 1750 base pairs, which was gel-purified. Half (by volume) of the purified product underwent CpG-methylation by *M.Sss*I methyltransferase supplemented with 640 µM S-adenosylmethionine (New England Biolabs) and purified using the QIAEX II kit (Qiagen). Completeness of methylation was checked by the inability to be digested by the methyl-sensitive *Hpa*II but successful digestion by the methyl-insensitive *Msp*I. Subsequently, in parallel, equal concentrations (∼100 ng/reaction) of the non-methylated and methylated fragments were treated with increasing levels of MNase (0U – 0.1U – 1U – 5U – 25 U) for 5 minutes at 22°C in 1X MNase digestion buffer. The MNase reaction was halted by addition of 6X purple loading dye (New England Biolabs) and electrophoresed immediately on a 1% agarose gel at 50 V for 20 minutes, followed by 100 V for 100 minutes. Gene Ruler Mix (Thermo Scientific) was loaded in the first lane. Electrophoresed products were stained with ethidium bromide, visualized by UV imaging (Fig. S4B, left hand side), and DNA fragments were transferred to a positively-charged nylon membrane, and probed with a 177 bp probe, annealing just upstream of the GGGGCC repeat tract (Table S1). To maximize our ability to assess that loss of signal corresponded to MNase digestion of the GGGGCC repeat fragment, we developed the 177 bp probe which anneals immediately adjacent to the repeat. (The 177 probe anneals 7 bp upstream of the GGGGCC repeat tract, while the 241 bp probe anneals 153 bp upstream). The 177 bp probe was produced by PCR using primers (Fwd: 5’–GAGCAGGTGTGGGTTTAGGA–3’ and Rev: 5’– CGACTCCTGAGTTCCAGAGC–3’) (Table S1). A phosphor image of the Southern blot was obtained using the Typhoon system (Fig. S4B, right hand side), and the rate of digestion of probed sequence was calculated using densitometric analysis in ImageJ. The percentage of DNA resistant to MNase (therefore, the fraction of undigested ‘starting fragment’ BAC DNA) was plotted as a function of MNase units of activity (concentration) in GraphPad Prism 8 (Fig. S4B).

### MNase controls

The MNase resistance we observe is not a reflection of the sequence preference of MNase or the sequence composition of the *C9orf72* region. The MNase resistance spanning the mutant *C9orf72* locus supports altered chromatin packaging of the non-repetitive flanks, as the 241 bp probe anneals 153 bp upstream of the (GGGCC)n repeat, and resistance is still evident (Fig. 5). The relative resistance of the expanded allele compared to the internal control of the non-mutant *C9orf72* allele, provides strong support for altered chromatin accessibility. MNase is known to display a cleavage sequence preference, preferring 5′-NpT and 5′-NpA sites (226, 227), yet is capable of digesting G-C only sequences and can digest most calf thymus DNA with over-digestion, which diminishes MNase cleavage preference(226). All of our nuclear digests were completed to over-digestion. The *C9orf72* sequences flanking the repeat, that are probed herein, contain an extensive number of the preferred 5′-NpT and 5′- NpA sites (Fig. S2B, green font). That protein-free genomic DNA from *C9orf72* expansion carriers is digested at an equal rate relative to the internal equimolar control of the non-expanded *C9orf72* allele, supports that the flanks can be digested (Fig. S4A). The MNase digest of the purified (GGGGCC)160 repeat DNA further confirms that the pure GC-rich repeat can be digested by MNase as evidenced by the ethidium bromide-stained gel (Fig. S4B). As such, the reduced digestion of the expanded allele in its native chromatinized state represents a chromatin-mediated inaccessibility.

To rule-out possible artefacts such as differences in the sensitivity between ethidium bromide staining and Southern probe hybridization, or loss of DNA during Southern transfer, or an uneven transfer or membrane retention of DNA, an aliquot of the same samples were run on a high-resolution gel, ethidium-stained, hybridized with the *C9orf72*-specific probe, imaged, and then the blot was stripped and re-probed using radiolabeled total genomic DNA as the probe (Fig. S4A). The hybridization pattern from total genomic DNA probe is very similar to the ethidium bromide-stained gel: most of the genome has been digested to completion to mono-, di-, tri-, and tetra-nucleosome sized fragments (Fig. S7A). In contrast, hybridization with the *C9orf72* probe shows dramatic MNase resistance of the expanded allele, while the non-expanded allele appears to be digested at the same rate as the rest of the genome.

Control MNase digestions of protein-free genomic DNA of unmethylated and methylated *C9orf72-*ALS samples, compared to the chromatinized samples of the same cell lines showed that MNase digestion occurs similarly for both alleles at extremely low MNase concentrations (0.01 Units) in protein-free conditions, yet the expanded allele is strongly resistant in chromatinized samples, supporting that this resistance is due to an unusual chromatin state (Fig. S4B-C).

There is a striking difference in MNase digestion between the ethidium-stained bulk genomic DNA, and the Southern blot detected *C9orf72* expanded allele, which is internally controlled for by its non-expanded homolog (Fig. 5). See above notes and Figure S4 for controls. The *C9orf72* region showed reduced presence in the mono-, di-, and tri-nucleosomal digested regions, relative to the rest of the genome (compare Southern blot with ethidium stain, Fig. 4A).

Control MNase digests using cloned *C9orf72* expanded (GGGGCC)160 repeat and flanks (+/-CpG-methylation) were performed to assess the impact of MNase sequence selectivity and the influence of CpG-methylation on digestibility. As shown in Figure S4B, the expanded repeat itself can be completely digested by MNase, and under the conditions used, the effect of sequence and CpG-methylation had limited effects on MNase digestibility, as previously reported (226, 227). This is consistent with the extensive number of the MNase- preferred 5′-NpT and 5′-NpA sites (226, 227) in the *C9orf72* sequences flanking the repeat, that are probed in the Southern blot (Fig. S2B, green font). These controls, coupled with the equal rate of MNase digestion of the protein-free *C9orf72* expanded allele in genomic DNA relative to the internal control of the non-expanded *C9orf72* allele (Fig. S4A-ii), demonstrates that the expanded repeat and flanks can be digested. Thus, the inaccessibility of MNase to the chromatinized *C9orf72* expanded allele and flanks (where the Southern probe binds), is due to its unusual chromatin compaction, and is not due to an inherent inability of MNase to cleave the expanded (GGGGCC)n DNA sequence.

Further mapping of the MNase accessibility in the chromatinized *C9orf72* expanded cells, with and without aberrant methylation, was performed. Essentially, for large expansions the mutant *C9orf72* allele is less accessible to MNase ∼3,300 bp upstream and ∼2,500 bp downstream of the expanded repeat. Decreased accessibility is further exacerbated by aberrant methylation (detailed in Fig. SD4A-iii). The absolute boundaries of decreased MNase accessibility varied with repeat expansions.

### Mapping MNase accessibility boundaries

We mapped the boundaries of MNase resistance in patient cell lines with no expansions (WT), expansions (P1, P7), and expansions with methylation (P3, P6), using a series of post-MNase restriction digests upstream and downstream of the repeat; fragments were detected by Southern blotting using either the 241 bp or 177 bp probe, as indicated. Upstream mapping was carried out with a post-MNase genomic digestion using *Ban*I, which cleaves just downstream of the repeat, in combination with either one of a series of restriction digests progressing along the upstream flank, as far as 3335bp upstream (*Afl*II, *Eco*RI, *Pvu*II, *Sph*I) (Fig. S4C). These samples were assessed by Southern blot, detected with the 177 bp probe (Fig. S4C upper panel, schematic). For each restriction fragment, the non-expanded allele signal was completely lost while the expanded allele was either resistant or partially digested to a shorter, distinct, nearly repeat-only containing fragment. This might suggest that the repeat tract itself is the most resistant to MNase accessibility. The amount of MNase-resistant starting material decreases progressively with increasing distance upstream from the repeat (Fig. S4C). Thus, the MNase resistance of the mutant allele extends upstream of the repeat at least ∼1252 bp to the *Pvu*II site, where some of the full-length fragment is still inaccessible to MNase, but diminishes further upstream ∼3335 bp to the *Sph*I site (Fig. S4C). Downstream mapping was conducted using a post-MNase genomic digestion with *Afl*II, which cleaves upstream of the repeat and beside the region detected by the 241 bp probe. *Afl*II digestion was paired with one of a series of restriction digests progressing along the downstream flank, as far as 3750 bp downstream (*Ban*I, *Bst*YI, *Bam*HI, *Pvu*II, or *Psi*I) (Fig. S4C lower panel, schematic). For each restriction fragment, the expanded allele was resistant and progressively digested to a shorter, distinct nearly repeat-only containing fragment, again suggesting that the repeat tract is the most MNase resistant region. Resistance in the downstream region was more evident for the methylated allele, and for larger expansions. For the larger methylated sample, P3 (∼3000 repeats), the MNase resistance of the mutant allele extends downstream of the repeat at least ∼2440bp to the *Pvu*II site, where some of the full-length fragment is still inaccessible to MNase, but diminishes further downstream ∼3750 bp to the *Psi*I site (Fig. S4C). For the shorter methylated expansion P6 (∼770 repeats), the MNase resistance of the mutant allele extends downstream of the repeat at least ∼1714bp to the *Bam*HI site, where some of the full-length fragment is still inaccessible to MNase, but diminishes further downstream ∼3750bp to the *Psi*I site (Fig. S4C). Thus, the MNase resistance of the mutant allele extends at least ∼2440 bp downstream from the repeat to the *Pvu*II site, where some of the full-length starting fragment is still resistant downstream ∼3,750 bp to the distal *Psi*I site (Fig. S4C). The methylated and expanded allele was more resistant than the non-methylated expanded allele. In summary, for large expansions some of the mutant *C9orf72* alleles can be resistant to MNase ∼1250 bp, diminishing through to ∼3300 bp upstream and downstream resistance is ∼2440 bp, diminishing through to ∼3700 bp away from the expanded repeat, and resistance is further exacerbated by aberrant methylation.

### Southern blot - *HTT*

Southern blotting for the *HTT* allele was followed as described (228), modifying the probe to be 226 bp, which anneals 143 bp upstream of the (CAG)n repeat tract, and used selected restriction-endonuclease digestions of genomic DNA – permitting sensitive detection of repeat size and length heterogeneity at molar levels. The 226 bp probe was produced by PCR using primers (Fwd: 5’-CTTGCTGTGTGAGGCAGAAC-3’ and Rev: 5’-CGCAGGTAAAAGCAGAACCT-3’). Molecular weight markers were run alongside the DNA (Gene Ruler Mix, Thermo Scientific, and Roche Molecular Weight Marker II).

### Chromatin accessibility *in cellulo* with MNase

An HD patient fibroblast cell line, with an *HTT* expansion of (CAG)21/180 (GM09197) was assessed for MNase accessibility at *HTT*. Cell pellets were washed in ice-cold 1X PBS, re-pelleted, and permeabilized by ice-cold NP-40 lysis buffer (10 mM Tris pH 7.4, 10 mM NaCl, 3 mM MgCl_2_, 0.5% NP-40, 0.15 mM spermine, 0.5 mM spermidine) for 5 min. on ice. Nuclei were collected at 200 *xg* (10 minutes, 4°C), washed in 1X MNase digestion buffer (10 X stock: 100 mM Tris pH 7.4, 150 mM NaCl, 600 mM KCl, 5 mM spermidine, 1.5 mM spermine), re-pelleted, and subsequently resuspended in 1X MNase digestion buffer supplemented with 1mM CaCl_2_. Treatment followed with MNase for 5 min. at 22°C, at differing concentrations (0.5-20U). All digestion reactions were stopped with an equal volume of *Stop* solution (0.625 mg/mL proteinase K, 20 mM EDTA, 20 mM EGTA, 1% SDS diluted in 1X MNase buffer) and incubated at 37°C for a minimum of 1 hour. Reaction conditions were optimized to ensure over-digestion by MNase, which diminishes MNase cleavage preference. DNA was purified by phenol/chloroform extraction, RNase A treatment, and ethanol precipitation. A small amount of purified DNA was assessed for completeness of MNase digestion, by its conversion to mono-, di-, tri-nucleosome sized fragments as resolved by electrophoresis stained with ethidium bromide and visualized by UV imaging. 12 ug of each DNA sample was then restriction digested to completion to release the repeat-containing fragment and flanking regions – this increases resolution by electrophoresis. Following neutral transfer, the *HTT* containing regions were detected by Southern blot as described above.

### RNA-seq library preparation, sequencing, and analysis

Total RNA from the lymphoblastoid cell lines was isolated using the hot TRIzol extraction protocol (229). Briefly, samples dissolved in TRIzol were incubated in a thermomixer set to 55°C and 1,000 rpm for 10 min. After cooling to ambient temperature, RNA was isolated by using TRIzol Reagent and the Direct-zol RNA MiniPrep Kit with DNase treatment according to manufacturer’s provided protocol (Zymo Research). RNA was quantitated using Qubit RNA BR Assay Kit (Thermo Fisher Scientific). Strand-specific, rRNA-depleted RNA-seq libraries were prepared using the KAPA Stranded RNA-seq Kit with RiboErase HMR (Kapa Biosystems) per manufacturer’s instructions, except for the use of custom Illumina-compatible index primers to allow multiplexing. Library size distribution was assessed using the High Sensitivity NGS Fragment Analysis Kit (DNF-747) on a Fragment Analyzer (Agilent). 2 × 150 paired-end sequencing was performed using an Illumina NextSeq 500. Reads were aligned to the genome assembly GRCh38 (hg38). Salmon was used for transcript expression quantification and differential gene expression analysis was performed using DESeq2(230, 231). A heat map was generated in R using the ggplot2 package. Graph was generated and statistical analysis was performed by using GraphPad Prism 8.

### Developing murine Tissue-Derived Cell Lines

Mouse tissues for cytogenetic analysis of chromosomes followed published protocols (137–140, 232–237). Mouse care, sacrifice, and tissue harvesting was done on the same day, including three wild type and three transgenic mice, all 5 months of age. Mice aged 5- months, when cell proliferation in the CNS and peripheral organs had ceased at 2-months of age (137–139). Ear, lung, tail, cerebrum, cerebellum, brain stem, kidney, liver, and spleen were harvested. A section of each tissue was manually macerated then treated with DMEM + Trypsin-EDTA for 1 hour in a 37°C incubator. Cells were collected, spun down, then seeded in a T25 flask with DMEM media supplemented with 20% FBS and 2 mM L-Glutamate. Two flasks were prepared: One with 1% Pen/Strep added, and one without. Once confluent, cells were seeded in a new T25 in DMEM (supplemented with 15% Fetal Bovine Serum, Penicillin/Streptomycin (100 IU/mL, 100 µg/mL), and 2 mM L-Glutamate to begin fragile site induction. To avoid cell type selection and cell type loss due to an inability to adapt to cell culture, harvested tissues were purposely cultured for very short terms following tissue harvesting. When possible, cells were grown to 80% confluency. Avoiding high cell densities, or long periods at confluency, has been shown to diminish FRAXA expression (234). Tissues were harvested and immediately made into cultures. These cultures were grown and induced to express the fragile sites and cytogenetically assessed. There was neither transit delays nor storage of cell lines until fragile sites were induced in metaphase. All procedures, including colony housing and tissue harvesting, were done in-house. Precautions were taken to avoid delays following tissue harvesting, tissue freezing, extended culturing and storage of cells, as these have been noted to diminish fragile site expression (234–237).

## ACKNOWLEDGEMENTS

This study is dedicated to the memory of Dr. Stephen T. Warren (1953-2021), a pioneer and advocate of Fragile X syndrome, a mentor, and friend. We thank Robert H. Brown for providing a YAC-*C9orf72* clone, John Volpe, Shamima Keka Islam for counting micronuclei, Lazar Joksimovic, and Raymond Wong for his expertise in the preparation of metaphase spreads. We thank Olivia Beck and Lieve Wiericx for sharing their expertise in cytogenetic fragile site recognition and Edwin Reyniers and Stephen W. Scherer, Jeff MacDonald, Mehdi Zarrei, Mehdi Layeghifard, and Bhooma Thiruvahindrapuram for help with the der-Chr9 data, and Michael Fenech for support regarding micronuclei, NBuds, and NBlebs.

## DATA AVAILABILITY

Datasets generated during the current study supporting its findings have been deposited or made available as indicated:

• RNASeq gene expression data - available on Gene Expression Ombinus (GEO), accession #GSE262674.

• Whole genome sequencing data in the form of a CRAM file is available upon request to the corresponding author.

## INCLUSION AND ETHICS STATEMENT

All collaborators of this study who have fulfilled the criteria for authorship required by *NAR Molecular Medicine* and Oxford University Press journals are included as authors. Each author’s contribution is detailed in the Author Contributions list. Expectations for collaborators were agreed upon ahead of the study’s initiation, and authorship placement was agreed upon by each collaborator. This study was conducted in collaboration with researchers and physicians from distinct cultural, ethnic, and racial backgrounds. The findings of this study apply to members of the population globally, as well as locally in the regions where patient recruitment occurred. The research does not result in stigmatization, incrimination, discrimination, or personal risk to participants who provided samples. Local and regional research publications relevant to our study was considered and included in our bibliography. The Hospital for Sick Children Research Ethics Board oversight and approval for the herein study was obtained. REB for human tissue, cell and animal studies: #1000056560.

## AUTHOR CONTRIBUTIONS

Conceptualization: CEP and RFK

Fig. 1: CEP

Fig. 2: ML, MDW, CEP

Fig. 3: MM, MHMS, CEP, PAD, GAR, EK, MZ

Fig. 4: MM, IvdW, SS, RFK, CEP

Fig. 5: MHMS, TKP, CEP (sample collection by LZ, JK, AA, PMcG, PMcK, JR)

Fig. 6: MM*, MHMS, NS, SL, KY, MM†, MK, RKCY, PMcK, JR, LS, MSS, CEP

Fig. 7: NS, BM (training by MM*, MHMS, TKP)

Table 1: MM*, MK, and CEP

Fig. S1: ML, MDW, CEP

Fig. S2: MM*, MZ, ER, MHMS Fig. S3: MM*, IvdW, SS, RFK Fig. S4: MHMS

Fig. S5: MM*

Fig. S6: MM*, MM†, SL, NS, KY, MHMS, CEP Fig. S7: TA, EAP, CHF

Fig. S8: RKCY, RFK, MM*, LS

Fig. S9: NS

Fig. S10: BM (training by MM*)

Table S1: MHMS

Table S2: LZ, JK, AA, PMcG, PMcK, JR, CEP Table S3: LS, MK, PMcK, RKCY, JR, CEP Table S4: MK, RKCY

BAC DNA for patient *C9orf72* provided by: CJ, PJdeJ ALS patient cells provided by: PAD, GAR

Autopsy tissues prepared by: LZ, PMK, JR

Intellectual input and experimental design: CEP, RFK, EC, RW, MDW, MSS, ER Supervision: CEP and RFK

Writing – original draft: MM*, CEP

Writing – review & editing: CEP, MHMS, NS, MSS, MM† All authors provided feedback on the manuscript.

MM* = Mila Mirceta

MM† = Mohiuddin Mohiuddin

## SUPPLEMENTARY INFORMATION

Supplementary information is available for this paper by request.

## MATERIALS AVAILABILITY

Correspondence and requests for materials should be addressed to Dr. Christopher E. Pearson (mail to). Mouse models presented in this study are commercially available from Jackson Laboratories (FVB/NJ-Tg(C9orf72)500Lpwr/J).

