## Extended Data and Figures for "*C9orf72* expansion creates the unstable folate-sensitive fragile site FRA9A"

### EXTENDED FIGURES AND TABLES

#### Figure S1

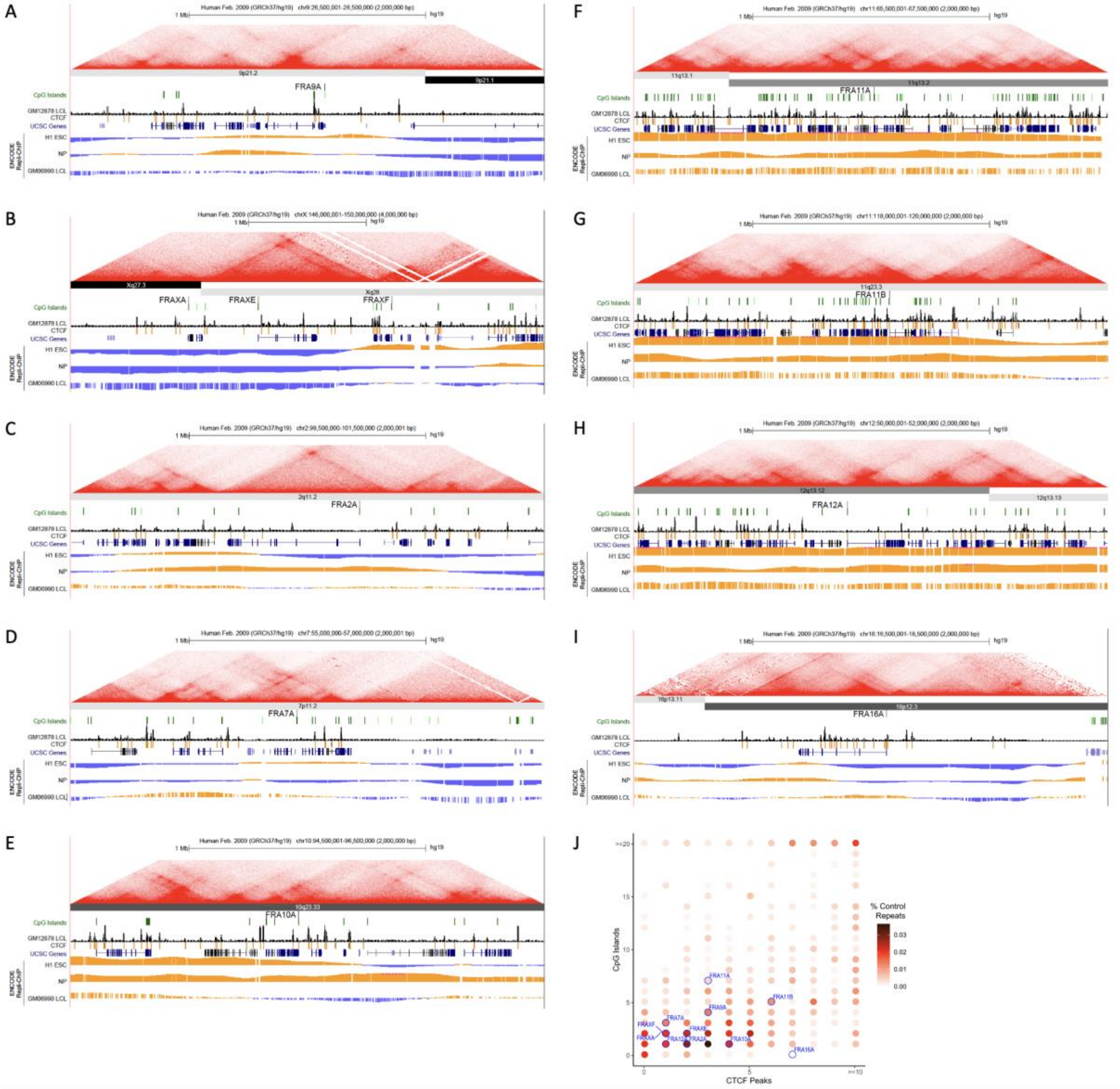

**Figure S1 Legend: The 9p21 locus and genomic features at FSFS and *C9orf72*.**

Topological associated domain (TAD), CpG island, and Repli-Chip analysis for all 10 CGG-folate-sensitive fragile sites FRA9A (A) FRAXA, FRAXE, and FRAXF (B), FRAq2A (C), FRA7A (D), FRA10A (E), FRA11A (F), FRA11B (G), FRA12A (H), FRA16A, and *C9orf72* (GGGGCC)n (FRA9A) (I). Shown are the contour density plots depicting the number of CTCF sites and CpG-islands in 100 kb windows centered on the repeat, representing boundaries with normal-length, matched repeats. Repli-seq data were obtained in bigwig format (GSE61972). H1 ESC = H1 Embryonic Stem Cells; accession:ENCFF000KUF; NP = Neuronal progenitor cells (BG01 Fibroblast-derived); accession:ENCFF907YXU; LCL = GM06990 lymphoblastoid cells; accession:ENCFF000KUA. Early (positive) and late (negative) replication was determined by subtracting the genome-wide median score from all values. To conserve space, only the FSFS gene is indicated, while other UCSC genes are condensed into one line. (J) Summary of enrichment of each fragile site (indicated in blue fonts) for CpG islands and CTCF sites. Points are colored according to density. See also Fig. 2.

.

**Figure S2**

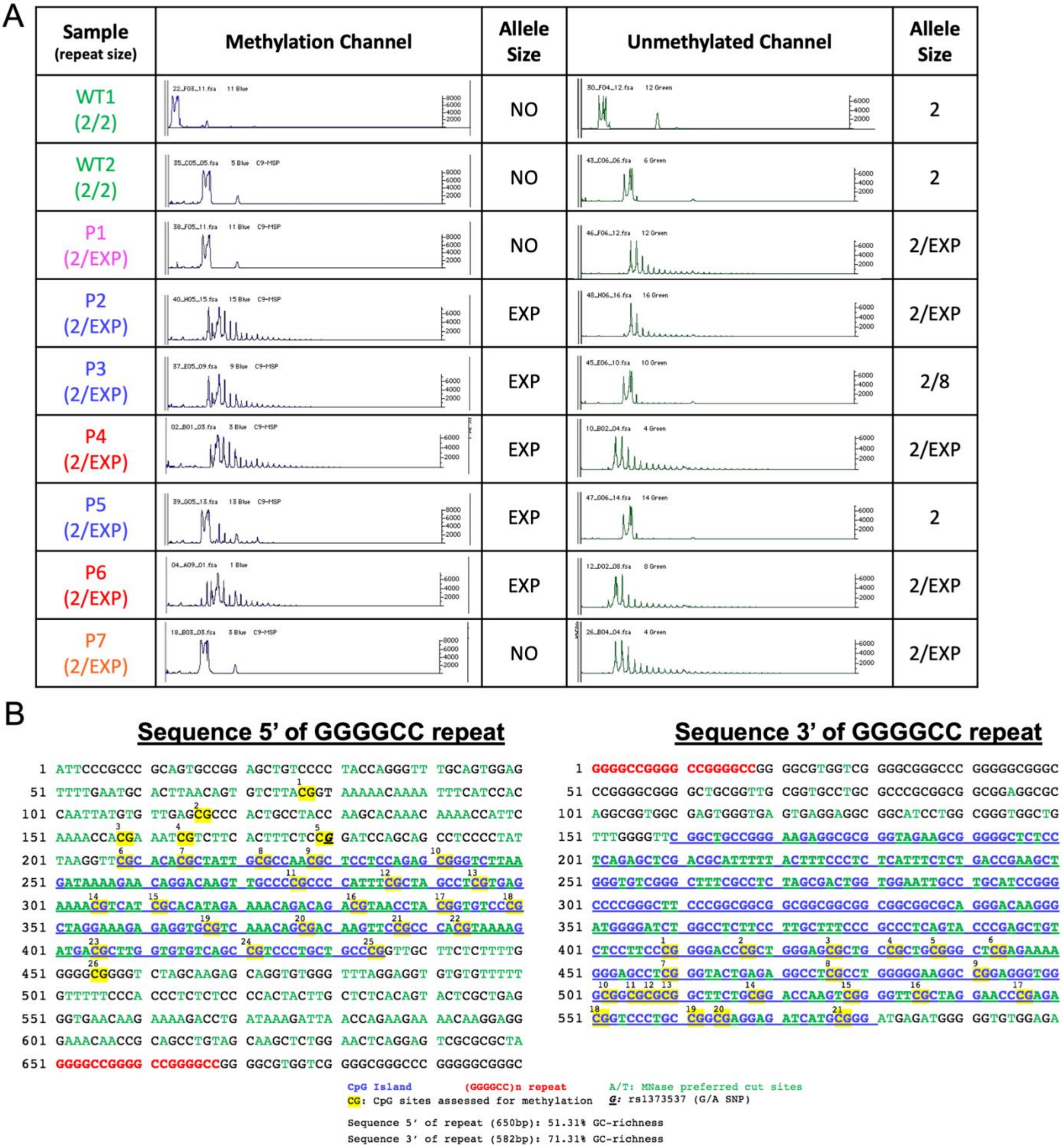
A
Sample (repeat size)
Methylation Channel
Allele Size
Unmethylated Channel
Allele Size
WT1 (2/2) | NO | 2
WT2 (2/2) | NO | 2
P1 (2/EXP) | NO | 2/EXP
P2 (2/EXP) | EXP | 2/EXP
P3 (2/EXP) | EXP | 2/8
P4 (2/EXP) | EXP | 2/EXP
P5 (2/EXP) | EXP | 2
P6 (2/EXP) | EXP | 2/EXP
P7 (2/EXP) | NO | 2/EXP
B
Sequence 5' of GGGGCC repeat
1 ATTCCCGCCC GCAGTGCCGG AGCTGTCCCC TACCAGGGTT TGCAGTGGAG
51 TTTTGAATGC ACTTAACAGT GTCTTACGGT AAAAACAAAA TTTCATCCAC
101 CAATTATGTG TTGAGCGCCC ACTGCCTACC AAGCACAAAC AAAACCATTC
151 AAAACCACGA AATCGTCTTC ACTTTCTCCG GATCCAGCAG CCTCCCCTAT
201 TAAGGTTCGC ACACGCTATT GCGCCAACGC TCCTCCAGAG CGGGTCTTAA
251 GATAAAAGAA CAGGACAAGT TGCCCCGCCC CATTTCGCTA GCCTCGTGAG
301 AAAACGTCAT CGCACATAGA AAACAGACAG ACGTAACCTA CGGTGTCCCG
351 CTAGGAAAGA GAGGTGCGTC AAACAGCGAC AAGTTCCGCC CACGTAAAAG
401 ATGACGCTTG GTGTGTCAGC CGTCCCTGCT GCCCGGTTGC TTCTCTTTTG
451 GGGGCGGGGT CTAGCAAGAG CAGGTGTGGG TTTAGGAGGT GTGTGTTTTT
501 GTTTTTCCCA CCCTCTCTCC CCACTACTTG CTCTCACAGT ACTCGCTGAG
551 GGTGAACAAG AAAAGACCTG ATAAAGATTA ACCAGAAGAA AACAAGGAGG
601 GAAACAACCG CAGCCTGTAG CAAGCTCTGG AACTCAGGAG TCGCGCGCTA
651 GGGGCCGGGG CCGGGGCCGG GGCGTGGTCG GGGCGGGCCC GGGGGCGGGC
Sequence 3' of GGGGCC repeat
1 GGGGCCGGGG CCGGGGCCGG GGCGTGGTCG GGGCGGGCCC GGGGGCGGGC
51 CCGGGGCGGG GCTGCGGTTG CGGTGCCTGC GCCCGCGGCG GCGGAGGCGC
101 AGGCGGTGGC GAGTGGGTGA GTGAGGAGGC GGCATCCTGG CGGGTGGCTG
151 TTTGGGGTTC GGCTGCCGGG AAGAGGCGCG GGTAGAAGCG GGGGCTCTCC
201 TCAGAGCTCG ACGCATTTTT ACTTTCCCTC TCATTTCTCT GACCGAAGCT
251 GGGTGTCGGG CTTTCGCCTC TAGCGACTGG TGGAATTGCC TGCATCCGGG
301 CCCCGGGCTT CCCGGCGGCG GCGGCGGCGG CGGCGGCGCA GGGACAAGGG
351 ATGGGGATCT GGCCTCTTCC TTGCTTTCCC GCCCTCAGTA CCCGAGCTGT
401 CTCCTTCCCG GGGACCCGCT GGGAGCGCTG CCGCTGCGGG CTCGAGAAAA
451 GGGAGCCTCG GGTACTGAGA GGCCTCGCCT GGGGGAAGGC CGGAGGGTGG
501 GCGGCGCGCG GCTTCTGCGG ACCAAGTCGG GGTTCGCTAG GAACCCGAGA
551 CGGTCCCTGC CGGCGAGGAG ATCATGCGGG ATGAGATGGG GGTGTGGAGA
CpG Island
(GGGGCC)n repeat
A/T: MNase preferred cut sites
CG: CpG sites assessed for methylation
G: rs1373537 (G/A SNP)
Sequence 5' of repeat (650bp): 51.31% GC-richness
Sequence 3' of repeat (582bp): 71.31% GC-richness

**Figure S2: Methylation analysis of *C9orf72*Exp patient cell lines.**

(A) Raw data of chromatograms for each patient analyzing repeat CpG-methylation as described (60). Data for methylated channel and unmethylated channel shown with column on right stating allele size observed in each channel. (B) Sequence upstream and downstream of the (GGGGCC)n repeat, including both the 5' and 3' CpG islands (blue). Bisulfite sequenced CpGs are highlighted in yellow and presence of >4 methylated sites was classified as a methylated CpG island. Both 5'NpA and 5'NpT sites are indicated in green font to show preferred micrococcal nuclease (MNase) digestion sites.

**Figure S3**

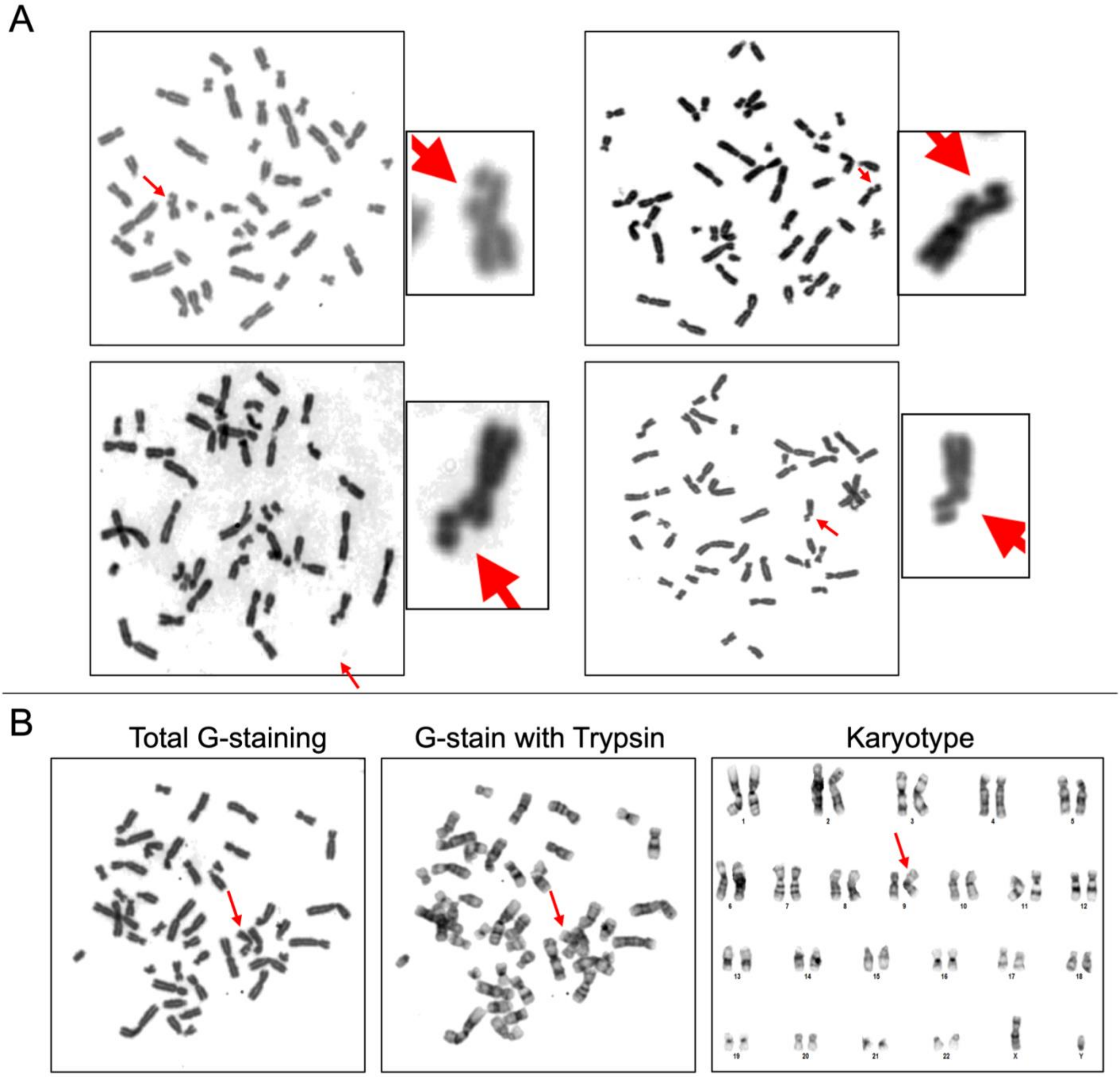

**Figure S3 Legend: Fragile site induction in folate-deficient conditions and Giemsa banding.** (A) Representative images of total Giemsa-stained metaphases with a fragile site induced in 199 media folate-deficient conditions. Red arrows point to fragile site. Magnified images to the right. (B) Example of metaphase with fragile site confirmed on Chr9 using total Giemsa stain, followed by trypsin treatment and karyotyping. Red arrows point to chromosome with fragile site.

**Figure S4**

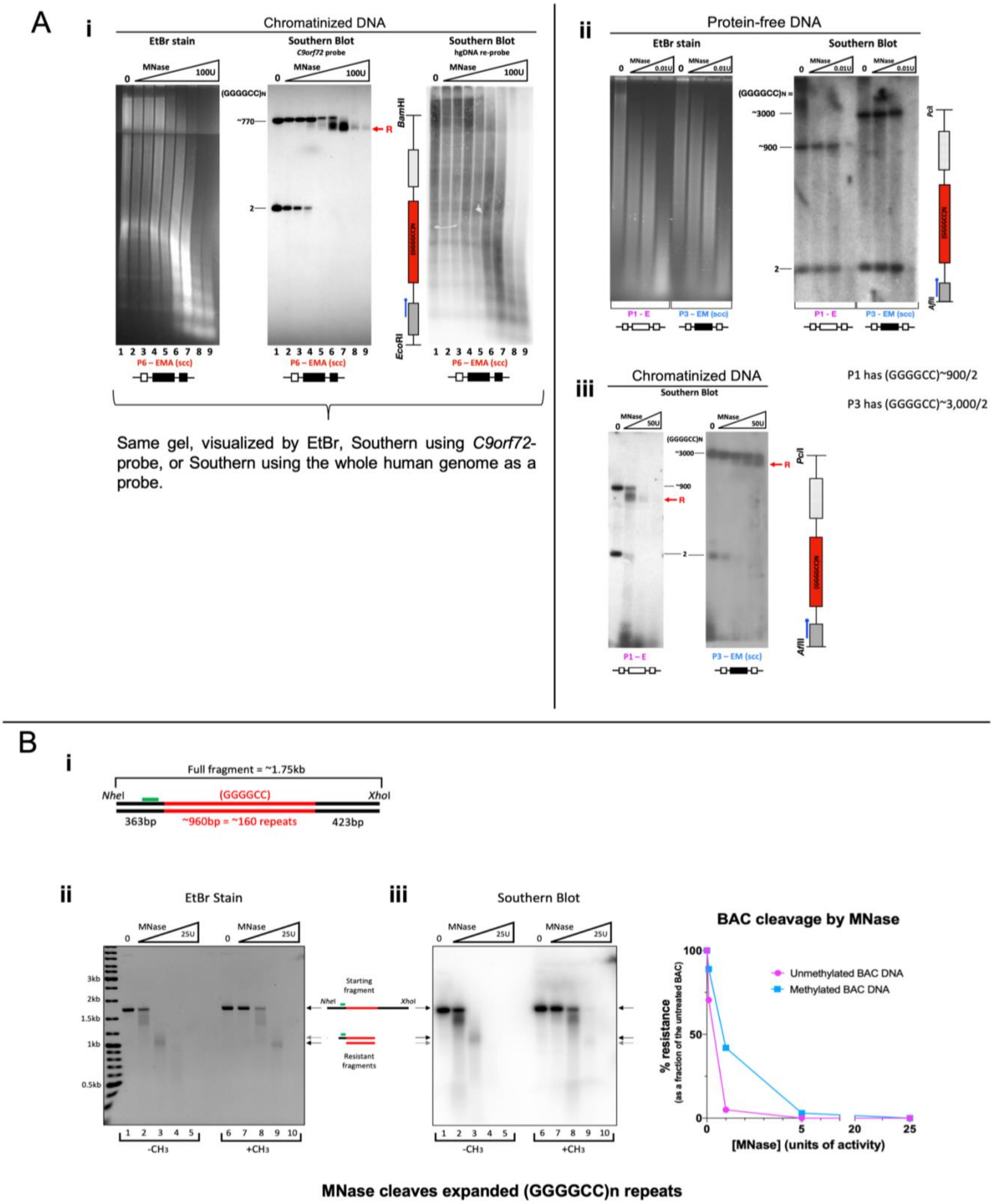

A
i
Chromatinized DNA
EtBr stain
Southern Blot
C9orf72 probe
Southern Blot
hgDNA re-probe
MNase
0
100U
(GGGGCC)N
~770
2
R
BamHI
EcoRI
1 2 3 4 5 6 7 8 9
P6 – EMA (scc)
Same gel, visualized by EtBr, Southern using C9orf72-probe, or Southern using the whole human genome as a probe.
ii
Protein-free DNA
EtBr stain
Southern Blot
0.01U
~3000
~900
2
P1 - E
P3 – EM (scc)
PciI
AflII
iii
Chromatinized DNA
Southern Blot
50U
P1 has (GGGGCC)~900/2
P3 has (GGGGCC)~3,000/2
B
i
Full fragment = ~1.75kb
NheI
(GGGGCC)
XhoI
363bp
~960bp = ~160 repeats
423bp
ii
EtBr Stain
25U
3kb
2kb
1.5kb
1kb
0.5kb
Starting fragment
Resistant fragments
1 2 3 4 5
6 7 8 9 10
-CH3
+CH3
iii
Southern Blot
BAC cleavage by MNase
Unmethylated BAC DNA
Methylated BAC DNA
% resistance
(as a fraction of the untreated BAC)
[MNase] (units of activity)
100
50
0
5
20
25
MNase cleaves expanded (GGGGCC)n repeats

**Figure S4 -continued-**

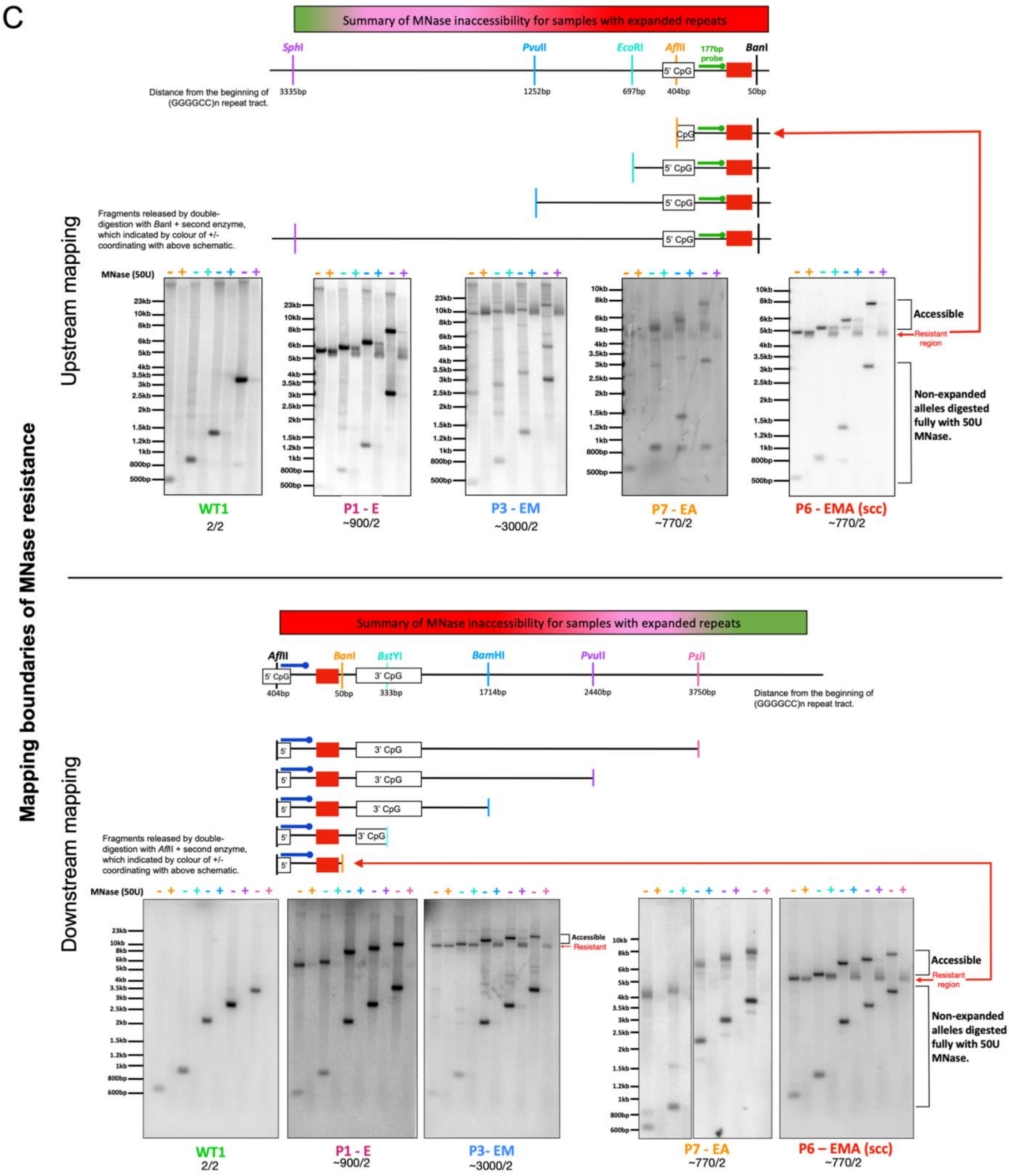

**Figure S4 Legend: MNase controls and MNase boundaries in *C9orf72*Exp cells.**

(A) MNase digestion of chromatinized cell DNA shows distinctly resistant bands in the expanded alleles, exclusive to chromatinized form of DNA. Related to Fig. 2. (Ai) Three images of the same gel: Ethidium bromide stained gel reveals completion of MNase digestion to nucleosomal fragments. Same gel was Southern blotted for *C9orf72*, showing MNase resistance of only the expanded allele. Membrane was stripped and re-probed using radio-labelled whole human genomic DNA; the whole genome is observed to undergo digestion at higher concentrations of MNase. Ultra-resolution of samples was attained by running them on a step-wise high resolution agarose gel (1.5% followed by 0.8%). (Aii & Aiii) MNase resistance depends upon chromatinization, as deproteinized “protein-free” *C9orf72*Exp gDNA (Aii) does not show differential MNase digestibility between the alleles. (Bi-iii) MNase can cleave the expanded (GGGCC)160 repeat +/-CpG methylation on a protein-free linearized bacterial artificial chromosome (BAC), where half was CpG-methylated by *M.SssI* methyltransferase. The repeat is digested and methylation has a measurable but minor effect on the ability of MNase to digest DNA. DNA was detected with the 177bp (green) probe (see Table ED5). The rate of digestion of the BAC starting fragment was calculated via densitometric analysis by Typhoon imaging and ImageJ. (C) Mapping boundaries of MNase-resistant region. Related to Fig. 2. Schematic, drawn to scale, reveals the *C9orf72* gene, gene motifs, probes and restriction sites used, where distance in base pairs upstream of the 5'-end of the GGGGCC repeats (upper panel) and base pairs downstream of the 3'-end of the GGGGCC repeats (lower panel). Upstream mapping involved MNase treatment, deproteinization, restriction digestions with *Ban*I and any one of the upstream restriction enzymes, followed by Southern blotting using the 177 bp green probe. Downstream mapping involved MNase treatment, deproteinization, restriction digestions with *Afl*II and any one of the downstream restriction enzymes, followed by Southern blotting using the blue probe. The repeat itself is fully resistant to MNase digestion, and the flanking regions adjacent to the repeat are significantly less accessible than the same regions on the nonexpanded allele (which is present at equimolar levels). Methylation appears to further enhance the observed resistance to MNase. MNase resistance diminishes as digests approach the *Sph*I site 3335 bp upstream of the repeat, while MNase resistance diminishes as digests approach the *Psi*I site 3750 bp downstream of the repeat.

**Figure S5**

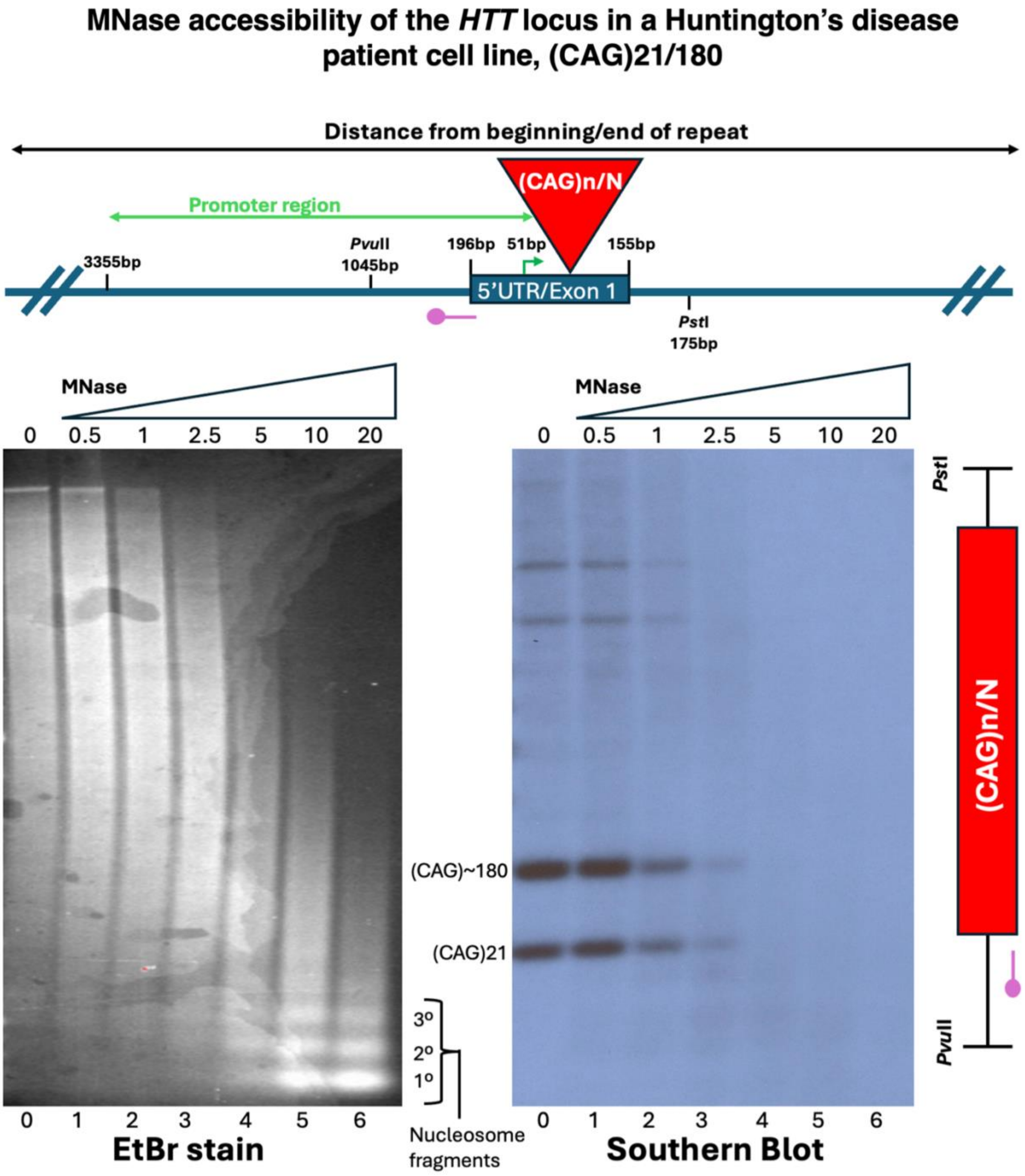

MNase accessibility of the *HTT* locus in a Huntington's disease patient cell line, (CAG)21/180
Distance from beginning/end of repeat
(CAG)n/N
Promoter region
3355bp
PvuII 1045bp
196bp
51bp
155bp
5'UTR/Exon 1
PstI 175bp
MNase
0 0.5 1 2.5 5 10 20
0 1 2 3 4 5 6
EtBr stain
(CAG)~180
(CAG)21
3º
2º
1º
Nucleosome fragments
MNase
0 0.5 1 2.5 5 10 20
0 1 2 3 4 5 6
Southern Blot
PstI
(CAG)n/N
PvuII

**Figure S5. MNase accessibility at the *HTT* locus with and without a CAG expansions.**
(A) *HTT* promoter/Exon 1 and restriction digestions and probe for Southern blot. MNase digestion of chromatinized cell DNA shows equally-accessible *HTT* alleles. Treatment was as outlined in Figure 5B and Methods.

**Figure S6**

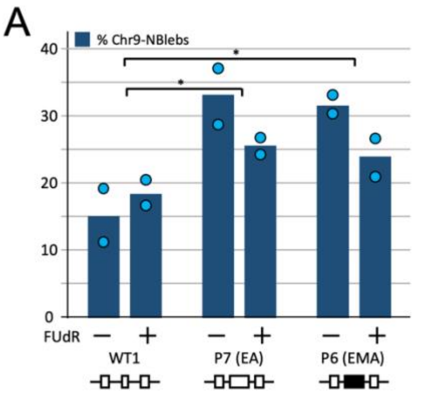


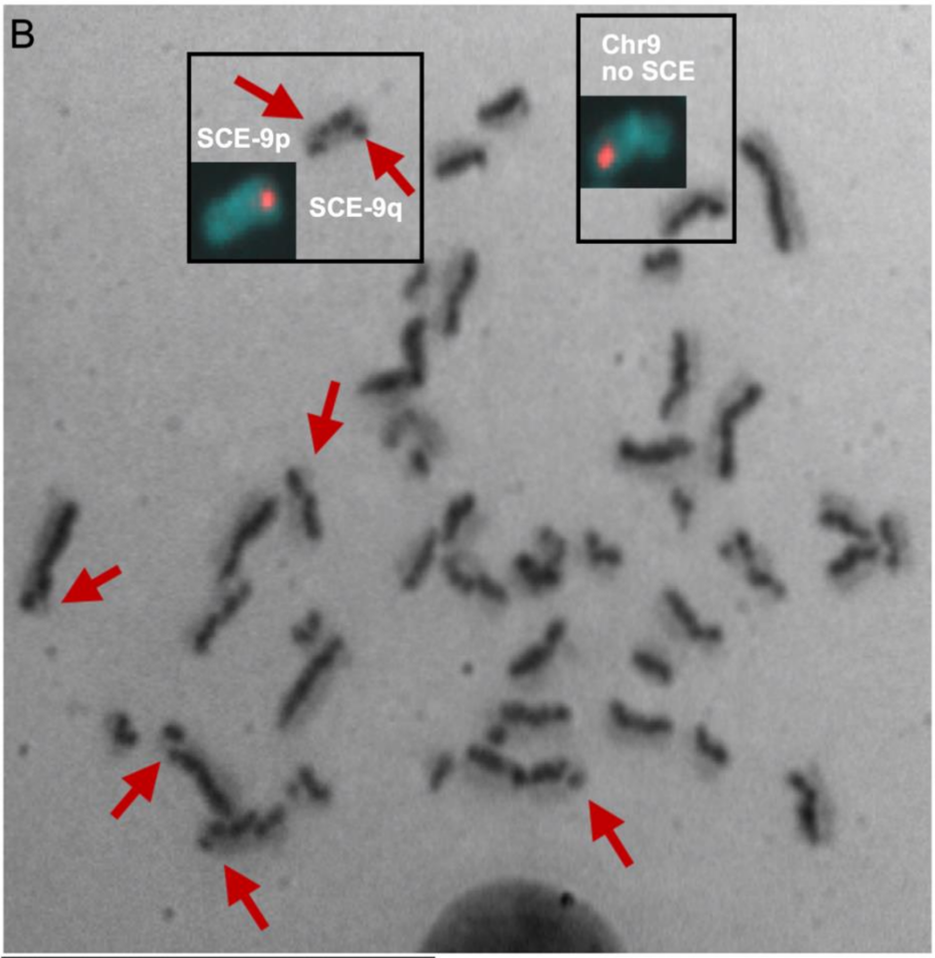


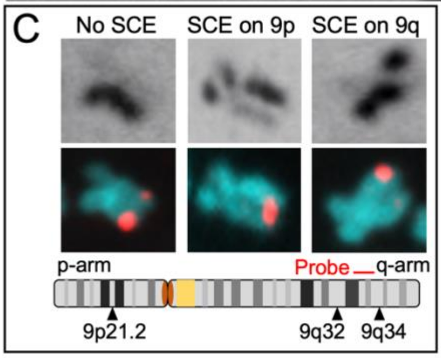


D

| | WT1 | P1 (E) | P5 (EM) | P6 (EMA) |
|---|---|---|---|---|
| No. of SCEs per chromosome 9[a] | | | | |
| Complete medium | 0.23 (17/37) | 0.26 (18/34) | 0.32 (22/34) | 0.37 (26/35) |
| Deficient medium | 0.23 (14/31) | 0.32 (25/39) | 0.35 (22/31) | 0.42 (32/38) |
| % of SCEs on chromosome 9[b] | | | | |
| Complete medium | 5.96 (17/285) | 5.66 (18/318) | 5.99 (22/367) | 5.65 (26/460) |
| Deficient medium | 5.88 (14/238) | 6.81 (25/367) | 6.67 (22/330) | 6.50 (32/492) |
| % of SCEs on Chr9p arm[c] | | | | |
| Complete medium | 35.3 (6/17) | 33.3 (6/18) | 31.8 (7/22) | 26.9 (7/26) |
| Deficient medium | 35.7 (5/14) | 56.0* (14/25) | 50.0** (11/22) | 46.9*** (15/32) |

Percent of sister chromatid exchange (SCE) at Chr9p for ALS patient cells grown in deficient medium are significantly different (red text) (asterisks refer to chi-square test above: *$\chi^2$= 10.43; df = 1; P <0.005, **$\chi^2$ = 6.85; df = 1; P <0.01 and ***$\chi^2$ = 8.67; df = 1; P <0.005)

[a]SCE calculations as per Wenger SL *et al.*, 1987, *Am J Med Genet*. **26**, 909-914. Data in parentheses are total number of SCE on Chr9 divided by total number of cells counted divided by 2 (2 Chr9s in each cell).

[b]Data in parentheses are the total number of SCE on chromosome 9 divided by total SCE on cells.

[c]Data in parentheses are total number of SCE on Chr9p divided by total SCE on Chr9.

**Figure S6. Nuclear blebs and SCEs in *C9orf72*Exp cells.**

(A) Quantification of nuclear blebs (NBlebs) in binucleated cells with *C9orf72* and 9q-FISH probes in WT, and *C9orf72*Exp cells (P6 (ssc) and P7) containing similar sized repeats (~770) but differing methylation status. Data are means of two independent experiments. Dots show data of each independent experiment. Statistical analyses using Fisher’s exact test (*P<0.05, **P<0.01, ***P<0.001). (B) Representative image of increased global expression of sister chromatid exchanges (SCEs) in *C9orf72*Exp cells, with Chr9-specific FISH images of Chr9 from the same spread overlaid on and beside the DAPI-stained Chr9s. (C) Representative images of Chr9 free of SCEs, with an SCE on 9p, or 9q, without SCEs on Chr9, or with SCEs on 9p or 9q. (D) Quantification of SCEs are significantly (Chi-square) increased at 9p relative to 9q in *C9orf72*Exp cells compared to WT cells.

**Figure S7**

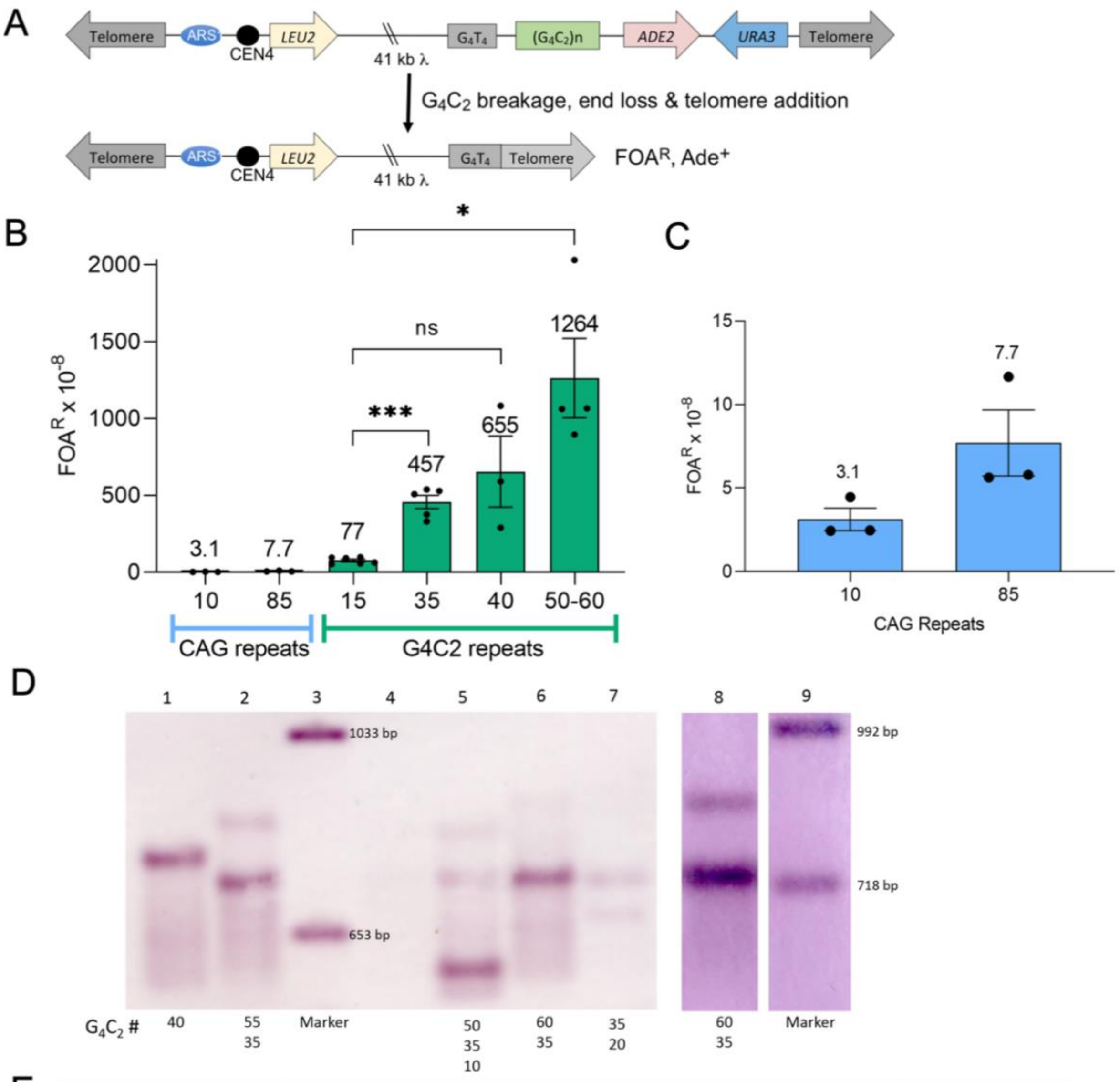


E

| Yeast strain # | Yeast strain name | Made from plasmid (sequenced repeat no. on plasmid) | $G_4C_2$ Repeat number on YAC, estimated by Southern blot | Number of $FOA^R$ colonies tested | Percent chromosome end loss |
|---|---|---|---|---|---|
| CFY 3234 | S3 | pCF504 (16 repeats) | ~15 repeats | 28 | 93% |
| CFY 3253 | M4-1 | pCF484 (29 repeats) | ~35, 55 repeats | 18 | 100% |
| CFY 3651 | M4-1a* | | ~10, 35, 50 repeats | 15 | 100% |
| CFY 3652 | M4-1b* | | ~20, 35, 60 repeats | 19 | 100% |
| CFY 3261 | M6-2 | pCF485 (36 repeats) | ~40 repeats | 22 | 100% |

*M4-1a and M4-1b are derivatives of M4-1.

**Figure S7. *C9orf72* GGGGCC repeat is hyper-prone to double-strand breakage in yeast.** (A) Repeat breakage as measured by yeast artificial chromosome (YAC) end loss. (B-G) End loss of chromosomes containing the indicated repeat number was measured by the rate of FOA-resistance (FOAR), indicating loss of the URA3 marker. Average measured rates are indicated above the bar for each strain; individual assays and SEM shown. (C) YAC end loss was confirmed by PCR and Southern blots. All colonies tested for end loss rate were also tested for starting (GGGGCC)n repeat length in the same experiment (see Methods). Note that "50-60" indicates the presence of lengths of 50-60 repeats but shorter repeat sizes were also present.

The YAC fragility assay detects chromosome end loss by selection for loss of the *URA3* gene (causing FOAR) and the *ADE2* gene (resulting in red colonies). Selection for Leu2+ ensures the broken chromosome is still present. This assay is an underestimate of the true DNA breakage rate, as it only measures events that result in end loss and healing by telomere addition at the telomere seed sequence (G4T4)13. (G) Results for YACs containing (CAG)10 or (CAG)85 repeats. The strains containing 50, 55, or 60 (GGGGCC)n repeats used all contained a substantial amount of shorter repeat heterogeneity, as indicated. Repeat tracts of 15, 35, and 40 (lane 1), were generally more homogenous in starting length, but with shorter sizes sometimes visible. Repeat sizes were determined by comparison to size standards and are approximate, rounded to the nearest five. Fragility Assay data for yeast strains containing the indicated YAC. *Rates are reported for colonies with the indicated repeat size; colonies with contracted or missing bands were not used in assays. Fragility assays using the M4 strains were split into those that contained a longer repeat species (50-60 repeat) and those that did not (e.g. primarily ~35 repeats). For colonies with 50-60 repeats, there was a mixed population with smaller sizes also present.

135 **Figure S8**

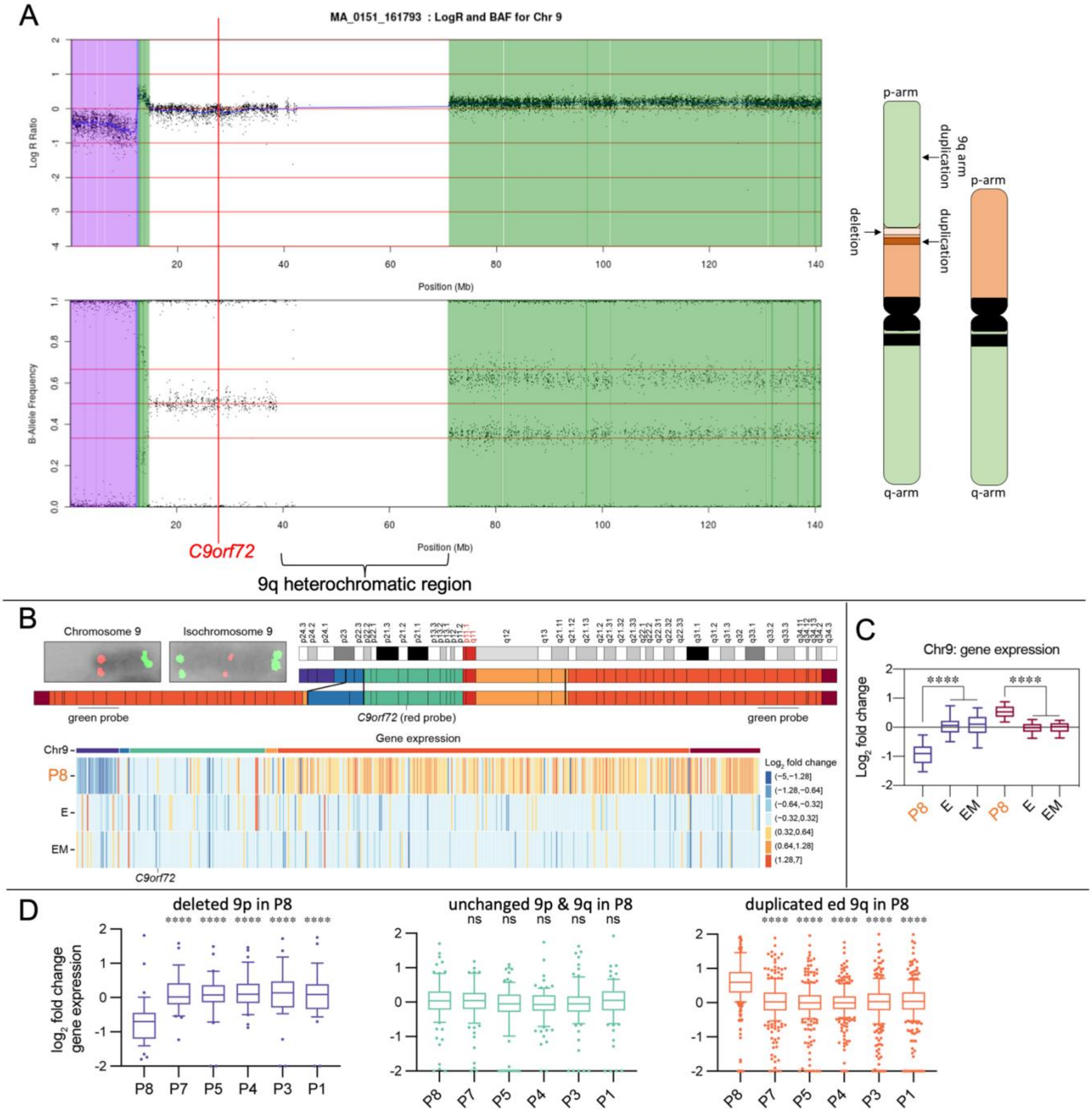

**Figure S8 Legend: CNV/SNP array and gene expression analysis of derivative-Chr9 of ALS-patient.** Related to Fig. 5. (A) Top illustrates probe intensity, bottom illustrates the B-allele frequency. A deletion of the short arm (arr 9p24.3p23(ter-12397876x1,12401666x2)) and a duplication of the long arm are evident (arr 9p23q34.3 (12687872x2,12709305-terx3)). *C9orf72* gene is indicated on the non-deleted or duplicated region in red. Rearrangement yields an isochromosome 9 [i(9q); del(9p)9p24-ter); dup(9q)9q21-ter] with a deletion of 9pter-p23 and a duplication of a short fragment of 9p23 and of the entire 9q arm was present. (B) Top: normal and rearranged chromosomes 9 ideograms. Positions of FISH green and red probes are indicated. Bottom: heat map shows highly expressed genes on chromosome 9 in the P8 *C9orf72*Exp-ALS cell line detected by RNA-seq, unmethylated (UM; n = 3), and methylated (M; n = 3) is normalized to unaffected controls (n = 4). (C) Box plot of gene expression changes in deleted (25 genes; violet), duplicated (282 genes), chromosome 9 regions. Centerline represents median and box-plot extends from 25th to 75th percentile, whiskers show 10-90 percentile. Significant difference was determined by Kruskal-Wallis test followed by Dunn's multiple comparisons test. **** for adjusted P-value $<0.0001$. (D) Previously published data for chromosomal instability.

**Figure S9**

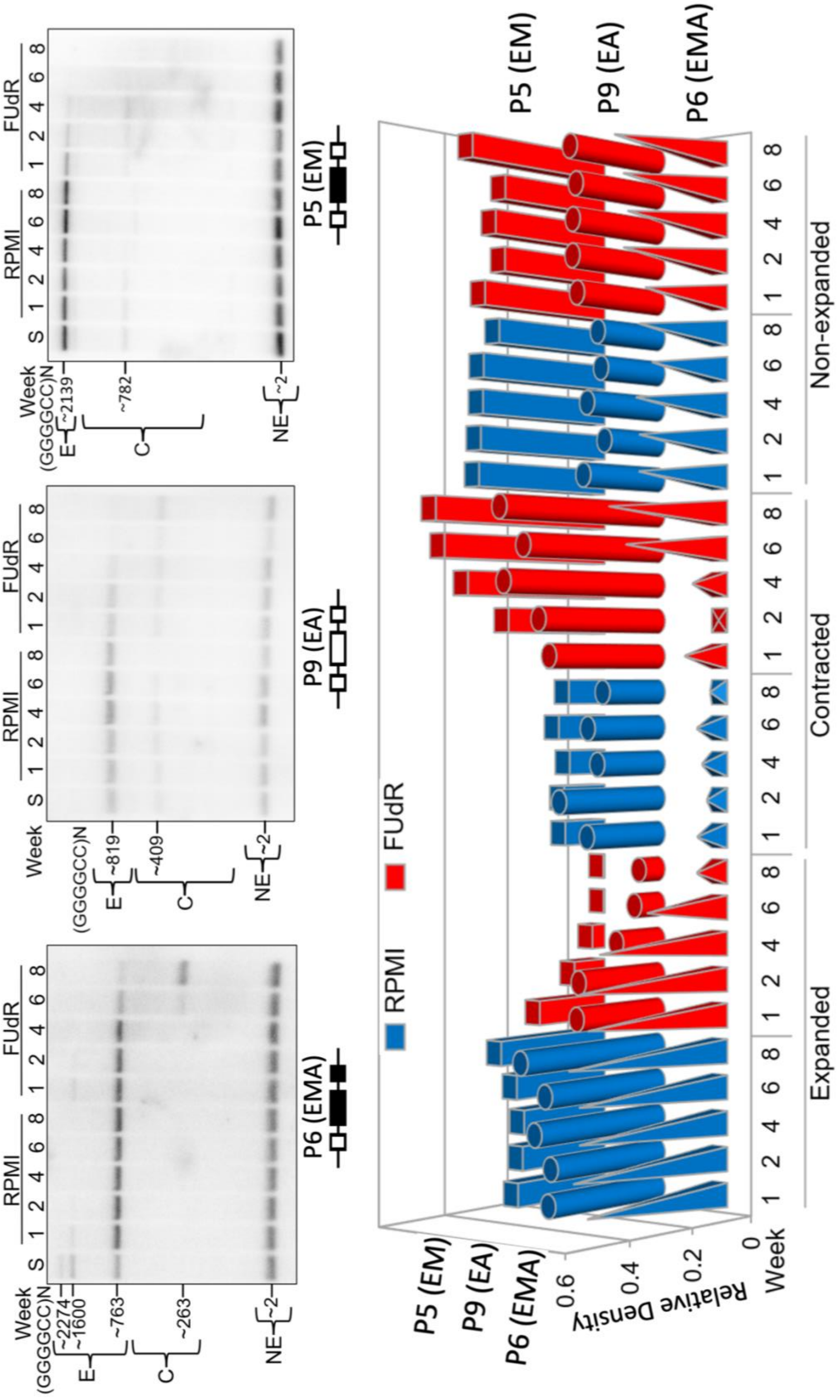

P6 (EMA)
P9 (EA)
P5 (EM)
RPMI
FUdR
Week
S
(GGGGCC)N
E
C
NE
~2274
~1600
~763
~263
~819
~409
~2139
~782
~2
Relative Density
Expanded
Contracted
Non-expanded

**Figure S9 Legend: *C9orf72*Exp is unstable during cell culture with FUdR.** *C9orf72*Exp carrying cells were grown for eight weeks in RPMI media or with addition of 0.1 µM FUdR. Southern blots run with DNA collected at indicated timepoints (Week). Changes in repeat size can be seen when comparing the expanded allele size at the start of culturing (S) to RPMI and RPMI with 0.1 µM FUdR throughout the eight weeks. Estimated repeat unit sizes, converted relative to a size marker, are indicated on the left of each blot ((GGGGCC)N). Results of all three Southern blots are represented in the graph (bottom). Signal densities for each band were calculated relative to the signal density of the entire lane (relative density) where a higher relative density corresponds to a larger cell population with a particular repeat size. The graph plots the relative density on the y-axis, the time-points (Week) grouped by repeat size (Expanded [E], Contracted [C], and Non-expanded [NE]) on the x-axis, and the three different cell lines (P6, P9, and P5) on the z-axis. RPMI conditions are plotted in blue, and FUdR conditions are plotted in red. The graph shows the decrease in cells with an expanded allele and an increase in cells with a contracted allele throughout the eight weeks upon FUdR treatment in comparison to RPMI conditions. The non-expanded allele remains stable throughout the 8 weeks in both conditions. No differences in trend are seen with differing methylation or clinical status (P6 – methylated, affected, P9 – no methylation, presymptomatic, P5 – methylated, presymptomatic). The effect we observed is similar to the recently reported effect of FUdR upon the somatic instability of the CGG repeat in FXS/FRAXA cells (98).

**Figure S10**

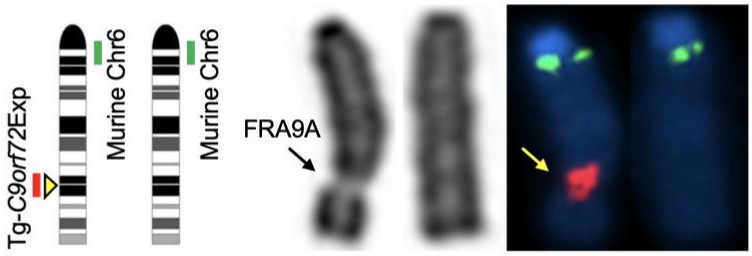


| Mouse | Fragility: % | Metaphase: n= | Repeat Size |
|---|---|---|---|
| Tg-1 | 15.5 | 100 | 185-680 |
| Tg-2 | 11.7 | 103 | 700-960 |
| Mean | 13.6 | | |
| WT-1 | 3.5 | 100 | |
| WT-2 | 1 | 100 | |
| Mean | 2.3 | | |

**Figure S10 Legend: *C9orf72* expanded transgene causes FRA9A in mouse tissues.** FRA9A expressed in fibroblasts (ear) of *C9orf72* transgenic mice, but not control mice (same magnification). Shown is a representative metaphase from cells prepared from one of three C9-500 mice, Tg-1 stained with Giemsa and human-specific *C9orf72*-containing (red = RP11-491J7, probe #4) and murine Chr6-specific probe (green = RP23-333A3). Table shows fragile sites from two transgenic mice.

**Table S1. PCR primers for Southern blot probes.**

**Table S1. Primers to generate Southern blot probes.**

| Probe | Primers Used | Figures Using This Probe |
|---|---|---|
| 241bp probe | Fwd: 5' – AGAACAGGACAAGTTGCCCC – 3' | Figure 1B; Figure 3C-F; Figure S4A & S4C, |
| | Rev: 5' – AACACACACCTCCTAAACCC – 3' | |
| 177bp probe | Fwd: 5' – GAGCAGGTGTGGGTTTAGGA – 3' | Figure S4C |
| | Rev: 5' – CGACTCCTGAGTTCCAGAGC – 3' | |
| Plasmid probe | *Psi*I – *Sac*I restriction fragment (943 bp) radio-labelled by random priming from ALS40 repeat plasmid | Figure S4B |

**Table S2 Summary of deep-phenotyping of PED25 family.**

**TABLE S2 - Deep-Phenotyping of PED25 family members Summary**



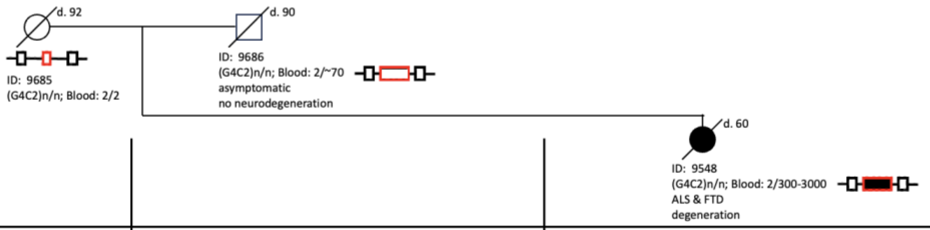


Xi Z P *et al.*, 2015, *AmJHumGenet*, **96**:962-70.

McGoldrick P *et al.*, 2018, *Neurology*, **90**: e323–e331.

Zhang MP *et al.*, 2017, *Acta Neuropathol.*, **134**: 271–279.

Clinical notes

| | | |
|---|---|---|
| **ID:**<br>**PED25, Canadian British family** | 9686 | 9548 |
| Age-at-onset (ALS, FTD):<br>Age-at-examination:<br>Age-at-death: | N/A (asymptomatic)<br>89-years<br>90-years | 57-years ALS & FTD<br>57-years<br>59-years |
| ***C9orf72* Risk-haplotype:** | Finnish haplotype | Finnish haplotype |
| **Genetic modifiers of *C9orf72*:**<br>*TMEM106B* rs1990622/rs3173615:<br>*ATXN2* CAG repeats: | AG/CG<br>(CAG)22/22. | AA/CC<br>(CAG)22/22. |
| **DNA methylation age-acceleration:** | DNAm age-acceleration = −8.8 years. | DNAm age-acceleration = +3.0 years |
| **(G4C2)2/N=:** | | |
| Blood: | 2/~70 | 2/~300-3000 |
| **CNS:** Frontal Cortex, Motor cortex, Occipital cortex, Temporal cortex, Hippocampus, Brainstem, Spinal cord (cervical, lumbar, & thoracic): | 2/~13-3500 (heterogenous, mostly 400-3000 repeats) | 2/~4400 (distinct size) |
| Cerebellum: | 2/~13-1500 (heterogenous, mostly 950-1500 repeats) | 2/~1600 (distinct size) |
| Heart, Tongue, Lung, Thalamus, Liver, Kidney, Epididymis, & Testicle: | 2/~13-3500 (heterogenous, mostly a distinct ~70 repeats) | 2/~4400 (distinct size) |
| **meCpG at *C9orf72*:**<br>Blood: | | |
| **CNS:** Frontal Cortex, Cerebellum, Motor cortex, Occipital cortex, Temporal cortex, Hippocampus, Brainstem, & Spinal cord (cervical, lumbar, & thoracic): | | |
| Liver, Kidney, & Skin: | | |
| Heart, Tongue, Lung, Thalamus, Epididymis, & Testicle: | | |
| Skin Fibroblasts: | | |
| **mRNA *C9orf72*:**<br>Blood:<br>Frontal Cortex:<br>Cerebellum: | Total *C9orf72* transcript levels from heart (2/70 alleles) and 5 mosaic tissues of 9686 were compared to the same tissues from 9548 (2/1,750 genotype). The mosaic carrier had higher *C9orf72* mRNA expression compared to 9548 (p < 0.05 for frontal cortex and p < 0.01 for other tissues) by 51% in cerebellum, 45% in frontal cortex. *C9orf72* mRNA expression levels in brain tissues (frontal cortex and cerebellum), cervical spinal cord and thoracic spinal cord, and non-CNS tissues (liver and heart) from 9686 (*C9orf72* genotype: 2/70/expansion) and 9548 (*C9orf72* genotype: 2/expansion). 9686 tissues carrying the small expansion had higher *C9orf72* expression levels than tissues with large expansions (*p < 0.05, **p < 0.01). A correlation analysis to understand the relationship between repeat expansion size and *C9orf72* expression. The ratio of Southern blot band intensity (70-repeat allele vs wildtype) in 9686 is significantly correlated with the ratio of *C9orf72* expression (9686 vs 9548) (p = 0.046). | |
| **C9ORF72 protein; C9-L & C9-S:**<br>Blood:<br>Frontal Cortex:<br>Cerebellum: | The long C9ORF72 Protein, C9-L (54 kDa) levels were present in the frontal cortex of both 9686 and 9548, and significantly higher in 9686 (*p = 0.044). The short C9ORF72 protein, C9-S (23 kDa) levels were similar between 9686 and 9548. C9-L was significantly increased in the urea fraction of cerebellum of 9686 compared to 9548 (*p = 0.042). C9-S levels in the HST fraction of cerebellum did not differ between 9686 and 9548. | |
| **RNA foci (sense & antisense):**<br>Blood:<br>Frontal Cortex:<br>Cerebellum: | Both sense-(GGGGCC) and antisense-(CCCCGG) RNA foci were evident in multiple brain regions of both father and daughter. Sense-(GGGGCC) RNA foci were evident in the frontal cortex (FCX), cerebellum (Purkinje and molecular cell layer), motor cortex (MCX), and hippocampus (HIP) of 9686 and 9548. Quantification of the total number of nuclei positive for sense RNA foci (I) revealed more (~2-fold) sense RNA foci–positive cells in the cerebellum of the unaffected 9686 compared to the ALS/FTD-affected daughter 9548 (n = 3, **p < 0.001); however, there were no differences in other regions. Antisense-(CCCCGG) RNA foci were also seen in the cerebellum, frontal cortex, motor cortex, and hippocampus of 9686 and 9548. Quantification of the number of antisense RNA foci–positive nuclei did not reveal any significant differences between 9686 and 9548. Similar levels of RNA-foci positive motor neurons in the spinal cords of 9686 and 9548. Quantification of sense- and antisense-RNA foci did not reveal a significant difference between 9686 and 9548 in cervical or lumbar spinal cord. | |
| **DPRs (GA, GP, GR, PA/PR):**<br>Blood:<br>Frontal Cortex:<br>Cerebellum: | All 5 DPRs were evident in both FCX and CBL of both father and daughter, consistent with the reported relative abundance of DPRs in *C9orf72* cases (GA > GP > GR > PA/PR). DPR burden in the granule cell layer of the cerebellum (CBL-GC) revealed significantly more (~7-fold) inclusions per field in the unaffected 9686 vs. the ALS/FTD-affected 9548: GA (205.8±11.62 vs. 30.33±4.13; ***p<0.001), GP (49.60±6.64 vs. 30.50±2.40; **p<0.01) and GR (7.48±0.66 vs. 1.00±0.22; ***p<0.001). DPR burden was similar in frontal cortex (FCX; M) and hippocampus (HIP; N). Although the number of GA-inclusions was significantly lower in motor cortex (MCX; O) of 9686 vs. 9548 (3.70±1.77 vs. 5.53±2.36; ***p<0.001), this was not found for other DPRs. | |
| **p62:** | Yes p62 inclusions,<br>See Neuropathology section. | Yes p62 inclusions,<br>See Neuropathology section. |
| **TDP-43:** | No TDP-43 inclusions,<br>See Neuropathology section. | Yes TDP-43 inclusions,<br>See Neuropathology section. |

Continued next page.

#### Table S2 -continued-

**TABLE S2 - Deep-Phenotyping of PED25 family members Summary (-continued-)** **Page 2 of 3**

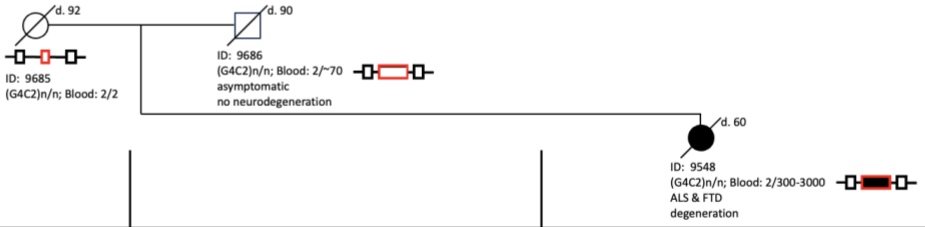


Xi Z P *et al.*, 2015, *AmJHumGenet*, **96**:962-70.

McGoldrick P *et al.*, 2018, *Neurology*, **90**: e323–e331.

Zhang MP *et al.*, 2017, *Acta Neuropathol.*, **134**: 271–279.

Clinical notes

| | | |
|---|---|---|
| **Neuropathology:** | No evidence of upper or lower motor neuron degeneration, and there was no TDP-43 pathology in the motor pathway. There was no evidence of FTLD TDP-43; the frontal and temporal cortices were not atrophic and did not contain TDP-43 inclusions. The most significant finding was the typical starburst-shaped p62 inclusions encountered in *C9orf72* carriers known to reflect DPRs. These inclusions were observed in the cerebellar granular layer, pyramidal neurons of CA1, and all sampled areas of the cerebral cortex. Other minor coexisting neurodegenerative phenomena included Alzheimer-type neuropathologic changes and scant TDP-43 pathology restricted to the mesial temporal regions (short neurites, neuronal cytoplasmic inclusions in the superficial layers of the entorhinal cortex, and dot-like inclusions of the dentate granular layer). Neuropathology of this 90-year old was not typical of presymptomatic *C9orf72* expansion carriers who can show pathology decades before onset (Bertrand A et al., 2018, *JAMA Neurol*, **75**:236-245; Querin G et al., 2019, *AnnNeurol*, **86**:158-167), suggesting that this individual truly asymptomatic. | At age 59 individual #9548 passed away and autopsy results confirmed the diagnosis of ALS with pathological signs of early stage FTLD. The neuropathologic diagnosis of the daughter (9548) was ALS with findings typical for *C9orf72* carriers. This included gliosis and loss of motor neurons from the anterior horn, TDP-43 and p62 inclusions in remaining motor neurons, and degeneration of corticospinal tracts. Neuronal p62 inclusions, some with a starburst morphology, were scattered throughout all cortical regions, the Ammon horn, the entorhinal cortex, and the cerebellar granular layer. There was a normal population of Betz cells, with some containing p62- and TDP-43–positive inclusions. |
| **Lifestyle:** tobacco: | Smoked two packs/day until his 60s, age that he started unknown. | Smoked a pack of cigarettes per day for 37 years starting at age 14 – 51 years.. |
| alcohol: | Previous alcoholic quit drinking when he was 32 then never drank again. | Drank a glass of red wine daily. |
| trauma/toxins: | No history of physical trauma, illegal drug abuse or pesticide exposure. | No history of physical trauma, illegal drug abuse or pesticide exposure. |
| Employment: | At 89-years he had been an officer in the British and Canadian navy, "always mentally sharp" as reported by his children. See Clinical section. | Had a college education and worked as an emergency unit clerk. See Clinical section. |
| **Clinical:** & other non-CNS presentations | At 89-years he had been an officer in the British and Canadian navy, "always mentally sharp" as reported by his children. At 89-years he was still independent for all activities of daily living. His performance fell within the normal range in the Social Norm Questionnaire with a score of 16/22, but he noted some difficulty with short-term memory over the last year (at age 89), which was reflected in a Montreal Cognitive Assessment (MoCA) score of 23/30, losing 2 points for mistakes with the clock drawing test and 5 points for delayed recall. Although the MoCA score of #9686 is within the range of mild cognitive impairment (18-26), it is still above the average (21) of the general elderly population of 70-80 years old. His mild cognitive impairment might indicate early disease symptoms; this MoCA score is still above the average for his age-group. Of note, 65% of Canadians older than 85 have cognitive impairment. He had a history of heart disease. At 90-years he passed away from late-onset chronic obstructive pulmonary disease (COPD). No cancers.<br><br>No evidence of corticobasal degeneration, behavioral variant (bvFTD), primary progressive aphasia (PPA), nonfluent/agrammatic variant (nfvPPA), apathy, socially inappropriate behavior, abnormal eating patterns, or loss of empathy, psychotic, hallucinatory, and/or delusional symptoms, visual illusions, strange ritual behaviors, bipolar disorder, schizophrenia, obsessive–compulsive disorder, and dementia with Lewy bodies, episodic memory issues emulating Alzheimer's disease, Parkinsonism with akinetic-rigid syndrome, or cerebellar signs.<br><br>No information on evidence of tremors, ataxia, myoclonus, dystonia, and/or chorea. | At 57-years of age developed symptoms of bulbar onset without any evidence of cognitive deficit on a Montreal Cognitive Assessment (MoCA) 3 (29/30). This individual had a college education and worked as an emergency unit clerk. The medical history included hypothyroidism, chronic obstructive pulmonary disease (COPD), cholecystectomy, incontinence, hysterectomy and a hernia. Individual #9548 Smoked a pack of cigarettes per day for 37 years starting at age 14 and drank a glass of red wine daily. At age 59 individual #9548 passed away and autopsy results confirmed the diagnosis of ALS with pathological signs of early stage FTLD.<br><br>Cholecystectomy, hernia and a hysterectomy in 2006 (purpose for hysterectomy unknown). No cancers.<br><br>No evidence of tremors, ataxia, myoclonus, dystonia, and/or chorea. No evidence of corticobasal degeneration, behavioral variant (bvFTD), primary progressive aphasia (PPA), nonfluent/agrammatic variant (nfvPPA), apathy, socially inappropriate behavior, abnormal eating patterns, or loss of empathy, psychotic, hallucinatory, and/or delusional symptoms, visual illusions, strange ritual behaviors, bipolar disorder, schizophrenia, obsessive–compulsive disorder, and dementia with Lewy bodies, episodic memory issues emulating Alzheimer's disease, Parkinsonism with akinetic-rigid syndrome, or cerebellar signs. |
| **Auto-immunity** [anti-DNA antibodies, lupus, psoriasis, skin sun-sensitivity, excessive warts, moles and/or lesions, skin redness (systemic lupus erythematosus or cutaneous lupus erythematosus), diabetes, arthritis, or multiple sclerosis]: | No evidence of autoimmune disorders. | Hypothyroidism. No other evidence of autoimmune disorders. |

Continued next page.

**Table S2 -continued-**

**TABLE S2 - NOTES & BIBLIOGRAPHY: (-continued-)** 

Characterization of the PED25 pedigree, including *C9orf72*Exp individuals that are asymptomatic and symptomatic, for clinical, lifestyle, neuropathology, and molecular biomarkers is described in detail (Xi Z *et al.*, 2015; McGoldrick P *et al.*, 2018; Zhang M *et al.*, 2017).

Related studies are also listed here.

Some factors could modify the *C9orf72*Exp phenotype. Intermediate *ATXN2* alleles (27–33 CAG repeats) render susceptibility to ALS and the *TMEM106B* A allele of rs1990622 was associated with a later onset ,while homozygosity for the G allele of rs3173615 protects against FTLD.9

Known *C9orf72* genetic modifiers (*TMEM106B* rs1990622, *TMEM106B* rs3173615 and *ATXN2* CAG repeats).

**Note:** Homozygosity for the minor allele (G) of rs3173615 in *TMEM106B* was reported to protect against developing FTD in *C9orf72*Exp carriers (Gallagher MD *et al.,* 2014, *Acta Neuropathol*, **127**:407-18). The major allele (A) of rs1990622 (conferring risk for developing FTLD) was associated to later age of onset and age of death in *C9orf72*Exp carriers (van Blitterswijk M *et al*., 2014, *Acta Neuropathol*, **127**:397-406).

Intermediate *ATXN2* alleles (27–33 CAG-repeats) were reported as modifiers in *C9orf72* carriers, rendering susceptibility to ALS (van Blitterswijk M *et al.,* 2014, *Neurobiol Aging*, **35**:2421.e13-7). These genetic modifiers were not able to explain the phenotype difference between 9548 and 9686.

**Penetrance:**

Recently the penetrance of the *C9orf72* expansion mutation has been reassessed and was found to be strikingly lower than initially thought: The mean risk of ALS at age 80 is 24%. but ranged between individual families from 16 to 60% (Van Wijk IF *et al.*, 2024, *AmyotrophLateralSclerFrontotemporalDegener*, **25**:188-196). The reduced penetrance and family-specific variations were suggested to be due to align "with the multistep hypothesis of ALS, suggesting the *C9orf72* repeat expansion, in isolation, is insufficient to cause ALS, but other genetic, epigenetic, or environmental factors are needed for the disease to manifest (Chiò A, *et al.,* 2018, *Neurology*, **91**: e635–e642).". Subsequent "steps" are unknown, but they likely determine disease penetrance, clinical variation, disease onset, progression and severity, all indispensable for family life decisions and clinical trial design.

Van Wijk IF *et al.,* 2024, Assessment of risk of ALS conferred by the GGGGCC hexanucleotide repeat expansion in C9orf72 among first-degree relatives of patients with ALS carrying the repeat expansion, *AmyotrophLateralSclerFrontotemporalDegener*, **25**:188-196.

Chiò A, *et al*., 2018, The multistep hypothesis of ALS revisited: The role of genetic mutations, *Neurology*, **91**: e635–e642.

**Genetic modifiers of *C9orf72*Exp:**

Zhang M, Tartaglia MC, Moreno D, Sato C, McKeever P, Weichert A, Keith J, Robertson J, Zinman L, Rogaeva E, 2017, DNA methylation age-acceleration is associated with disease duration and age at onset in *C9orf72* patients, *Acta Neuropathol*, **134**:271-279.

Gallagher MD *et al.*, 2014, TMEM106B is a genetic modifier of frontotemporal lobar degeneration with *C9orf72* hexanucleotide repeat expansions, *Acta Neuropathol*, **127**:407-18.

van Blitterswijk M *et al.*, 2014, TMEM106B protects C9ORF72 expansion carriers against frontotemporal dementia, *Acta Neuropathol*, **127**:397-406.

van Blitterswijk M *et al.*, 2014, Ataxin-2 as potential disease modifier in C9ORF72 expansion carriers, *Neurobiol Aging*, **35**:2421.e13-7.

**Neuropathology and presymptomatic *C9orf72*Exp:**

A long presymptomatic phase with evident degenerative changes are often detected decades before ALS or FTD symptom manifestation (Bertrand A et al., 2018, *JAMA Neurol*, **75**:236-245; Querin G *et al.,* 2019, *AnnNeurol*, **86**:158-167), pathology that was not present in the brain of the asymptomatic 90-year old father (McGoldrick P *et al.,* 2018, *Neurology*, **90**: e323–e331).

**Lifestyle (smoking, alcohol, diet, exposures, trauma, etc) & *C9orf72*Exp:**

Westeneng HJ, *et al.*, 2021, Associations between lifestyle and amyotrophic lateral sclerosis stratified by *C9orf72* genotype: a longitudinal, population-based, case-control study, *Lancet Neurol*, **20**:373-384.

**Cancer & *C9orf72*Exp:**

Tábuas-Pereira M *et al.*, 2019, Increased risk of melanoma in C9ORF72 repeat expansion carriers: A case-control study, *Muscle Nerve*, **59**:362-365.

**Autoimmunity & *C9orf72*:**

Béland LC *et al.*, 2020, Immunity in amyotrophic lateral sclerosis: blurred lines between excessive inflammation and inefficient immune responses, *Brain Commun,* 2:fcaa124.

McCauley ME *et al.*, 2020, *C9orf72* in myeloid cells suppresses STING-induced inflammation, *Nature*, **585**:96-101.

Burberry A *et al.*, 2020, *C9orf72* suppresses systemic and neural inflammation induced by gut bacteria, *Nature*, **582**:89-94.

Fredi M *et al.*, 2019, *C9orf72* Intermediate Alleles in Patients with Amyotrophic Lateral Sclerosis, Systemic Lupus Erythematosus, and Rheumatoid Arthritis, *Neuromolecular Med*, **21**:150-159.

Biasiotto G, Zanella I, 2019, The effect of *C9orf72* intermediate repeat expansions in neurodegenerative and autoimmune diseases, *Mult Scler Relat Disord*, **27**:42-43.

Chiò A *et al.*, 2018, The multistep hypothesis of ALS revisited: The role of genetic mutations, *Neurology*, **91**:e635-e642.

Burberry *et al.*, 2016, Loss-of-function mutations in the C9ORF72 mouse ortholog cause fatal autoimmune disease, *Sci Transl Med*, **8**:347ra93.

Continued next page.

**Table S3: Breakends and copy number variations in FRA9A-expressing derChr9 (P8) sample.**

→ *See coordinating Excel file*

**Table S4: Significant differentially expressed genes (DEGs) identified between wild-type and FRA9A-expressing derChr9 (P8) samples.**

→ *See coordinating Excel file*

#### AUTHOR CONTRIBUTIONS

Conceptualization: CEP and RFK

Fig. S1: ML, MDW, CEP

Fig. S2: MM*, MZ, ER, MHMS

Fig. S3: MM*, IvdW, SS, RFK

Fig. S4: MHMS

Fig. S5: MM*

Fig. S6: MM*, MM†, SL, NS, KY, MHMS, CEP

Fig. S7: TA, EAP, CHF

Fig. S8: RKCY, RFK, MM*, LS

Fig. S9: NS

Fig. S10: BM (training by MM*)

Table S1: MHMS

Table S2: LZ, JK, AA, PMcG, PMcK, JR, CEP

Table S3: LS, MK, PMcK, RKCY, JR, CEP

Table S4: MK, RKCY

#### MATERIALS AVAILABILITY
